# A unique sulfotransferase-involving strigolactone biosynthetic route in Sorghum

**DOI:** 10.1101/2021.09.08.459372

**Authors:** Sheng Wu, Yanran Li

## Abstract

LOW GERMINATION STIMULANT 1 (LGS1) plays an important role in strigolactones (SLs) biosynthesis and *Striga* resistance in sorghum but the catalytic function remains unclear. Using the recently developed SL-producing microbial consortia, we examined the activities of sorghum MAX1 analogs and LGS1. Surprisingly, SbMAX1d (accession # XP_002458367) synthesized 18-hydroxy-carlactonoic acid (18-hydroxy-CLA) directly from carlactone (CL) through four-step oxidations, and addition of LGS1 led to the synthesis of both 5-deoxystrigol (5DS) and 4-deoxyorobanchol (4DO). Further biochemical characterization found that LGS1 functions after SbMAX1d by converting 18-hydroxy-CLA to 18-sulphate-CLA to provide an easier leaving group to afford a spontaneous formation of 5DS and 4DO. The unique functions of SbMAX1 and LGS1 imply a previously unknown synthetic route towards strigolactones.

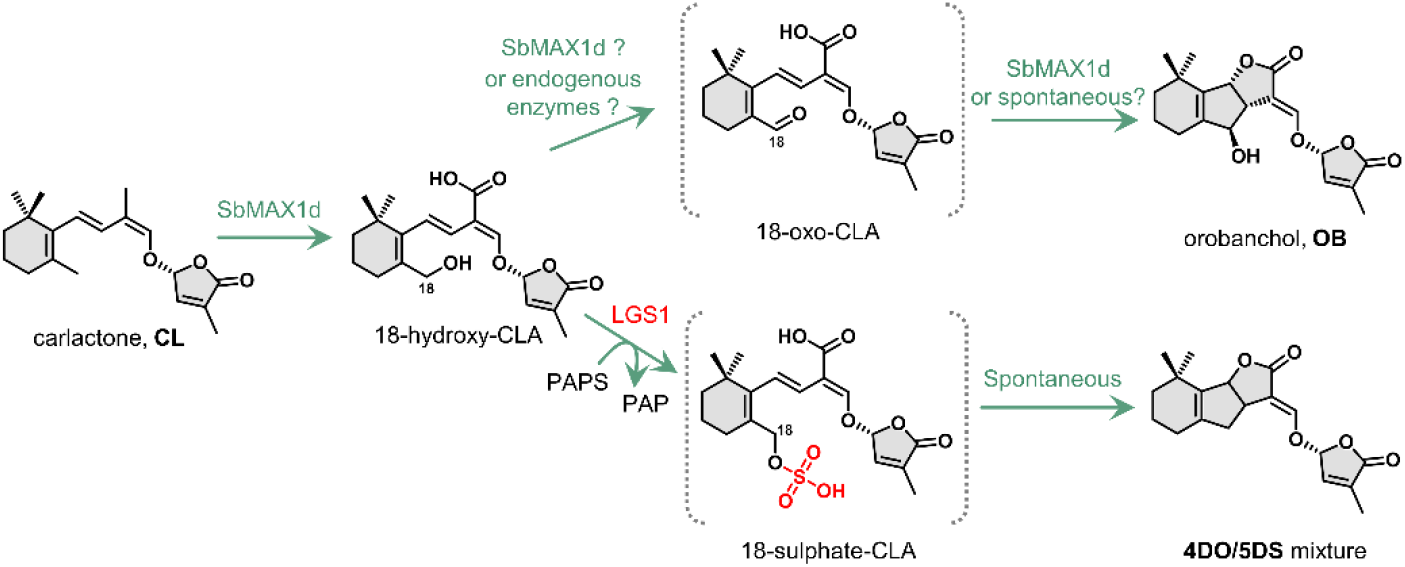

## Introduction

Strigolactones (SL) are a group of butanolide-containing molecules originally identified as seed germination stimulants for the parasitic weeds *Striga* and *Orobanche*^1, 2^, and later characterized as phytohormones that play diverse important roles in plant growth and development^3–5^. SLs can be divided into canonical and non-canonical SLs, with canonical SLs further grouped into strigol (*S*)- and orobanchol (*O*)-type SLs according to the stereochemistry of the C-ring^3^ (Figure 1). Different SL structures have been reported to exhibit distinct parasitic weed germination activities^6, 7^. For example, SLs exhibiting high germination stimulation activity towards *S. gesnerioides* induced low germination in *S. hermonthica*, while several SLs of high germination stimulation activity to *S. hermonthica* inhibit the germination of *S. gesnerioides*^8^. Recently, LOW GERMINATION STIMULANT 1 (LGS1) has been identified to be responsible for the *Striga* germination stimulant activity in sorghum and missing from the *Striga*-resistant sorghum varieties^9^, which produce distinct SL profiles, i.e., (*S*)-type 5DS and (*O*)-type orobanchol (OB), respectively^9^. LGS1 is a putative sulfotransferase (SOT), which normally catalyzes the transfer of a sulfonate group from 3′-phosphoadenosine 5′-phosphosulfate (PAPS) to a hydroxyl group of acceptor molecules^10^. The mechanism on how LGS1 regulates SL profiles between 5DS and OB in sorghum remains unclear.

**Figure 1.**
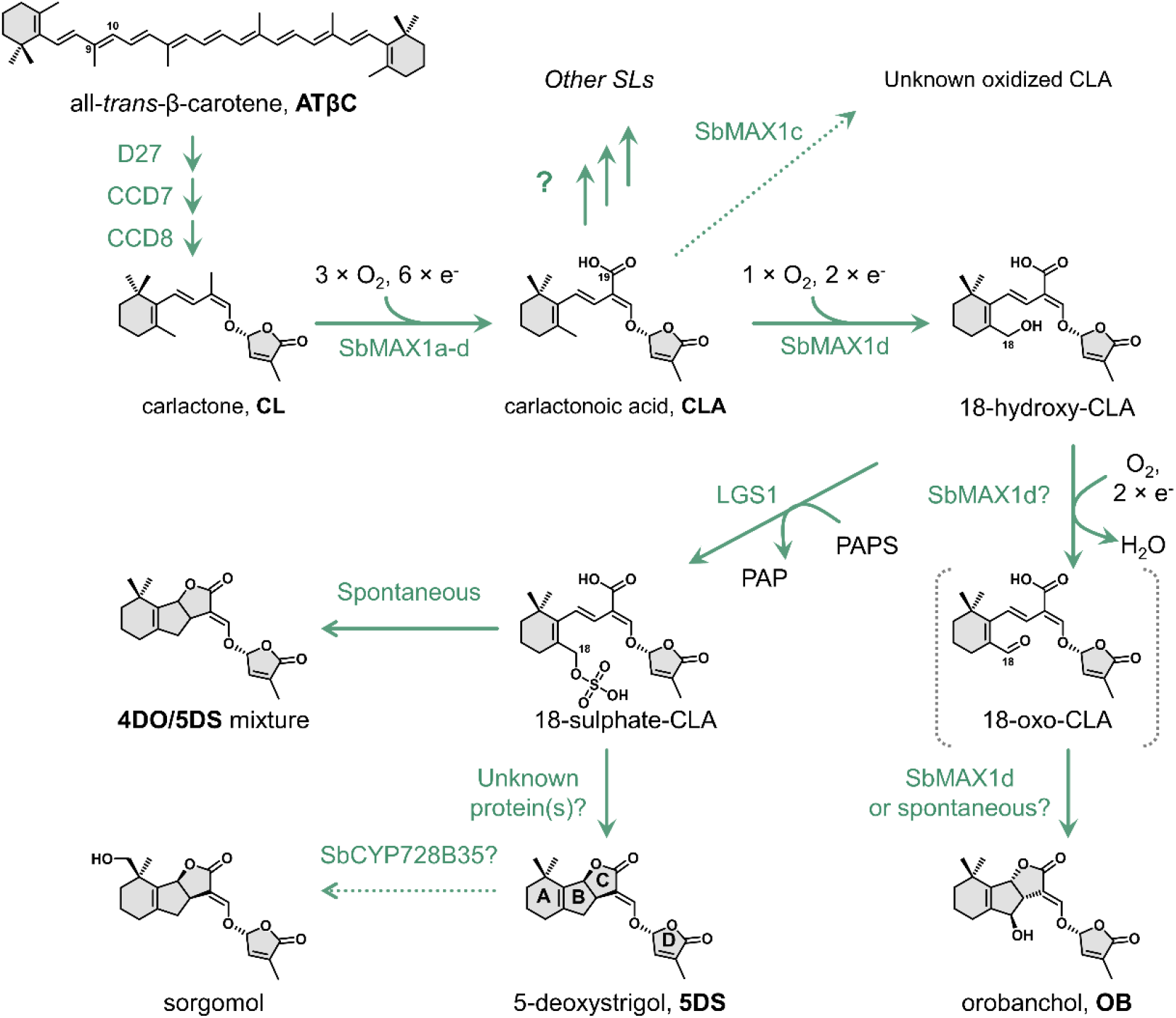
The proposed biosynthetic pathway of 5DS and OB in *Sorghum bicolor*. D27, [2Fe-2S]-containing isomerase DWARF27; CCD7, carotenoid cleavage dioxygenase 7; CCD8, carotenoid cleavage dioxygenase 8; SbMAX1d, MAX1 analog d from *S. bicolor*; LGS1, *LOW GERMINATION STIMULANT 1*, a sulfotransferase; PAPS, 3′-phosphoadenosine 5′-phosphosulfate; PAP, 3′-phosphoadenosine-5′-phosphate; 4DO, 4-deoxyorobanchol.

SLs are synthesized from carlactone (CL), which is then converted to diverse SL structures by various downstream tailoring enzymes especially cytochrome P450s (CYPs, Figure 1)^4, 11^. The two major groups of CYP that contribute to the structural diversity downstream of CL belong to CYP711A and CYP722C subfamily^12^. The best studied CYP711A is MORE AXILLARY GROWTH1 (MAX1) from *Arabidopsis thaliana* (AtMAX1), which converts CL to carlactonoic acid (CLA) and is functionally conserved in dicots^13^. On the other hand, monocots, especially the economically significant *Poaceae* family, often encode more than one CYP711As (Table S1, Figure 2a, S1), with diverse functions distinct from AtMAX1^13–16^. For example, rice has five MAX1 homologs, with CYP711A2 catalyzing the conversion of CL to 4-deoxyorobanchol (4DO) and CYP711A3 further oxidizing 4DO to OB^15^. Most CYP711As encoded by monocot plants remain to be characterized. The other major group of SL-synthesizing CYPs, CYP722C subfamily, catalyze the conversion of CLA towards either OB or 5DS^17–19^. Currently, there are two known routes towards the synthesis of (*O*)-type SLs catalyzed by either Group I CYP722C (e.g., VuCYP722C) or OsCYP711A2^15, 18^, while the only known 5DS biosynthetic route is through Group II CYP722C (e.g., GaCYP722C)^17^. However, CYP722Cs are generally missing from the *Poaceae* family including sorghum, which implies that sorghum employs a previously unknown strategy to synthesize (*S*)-type SL.

**Figure 2.**
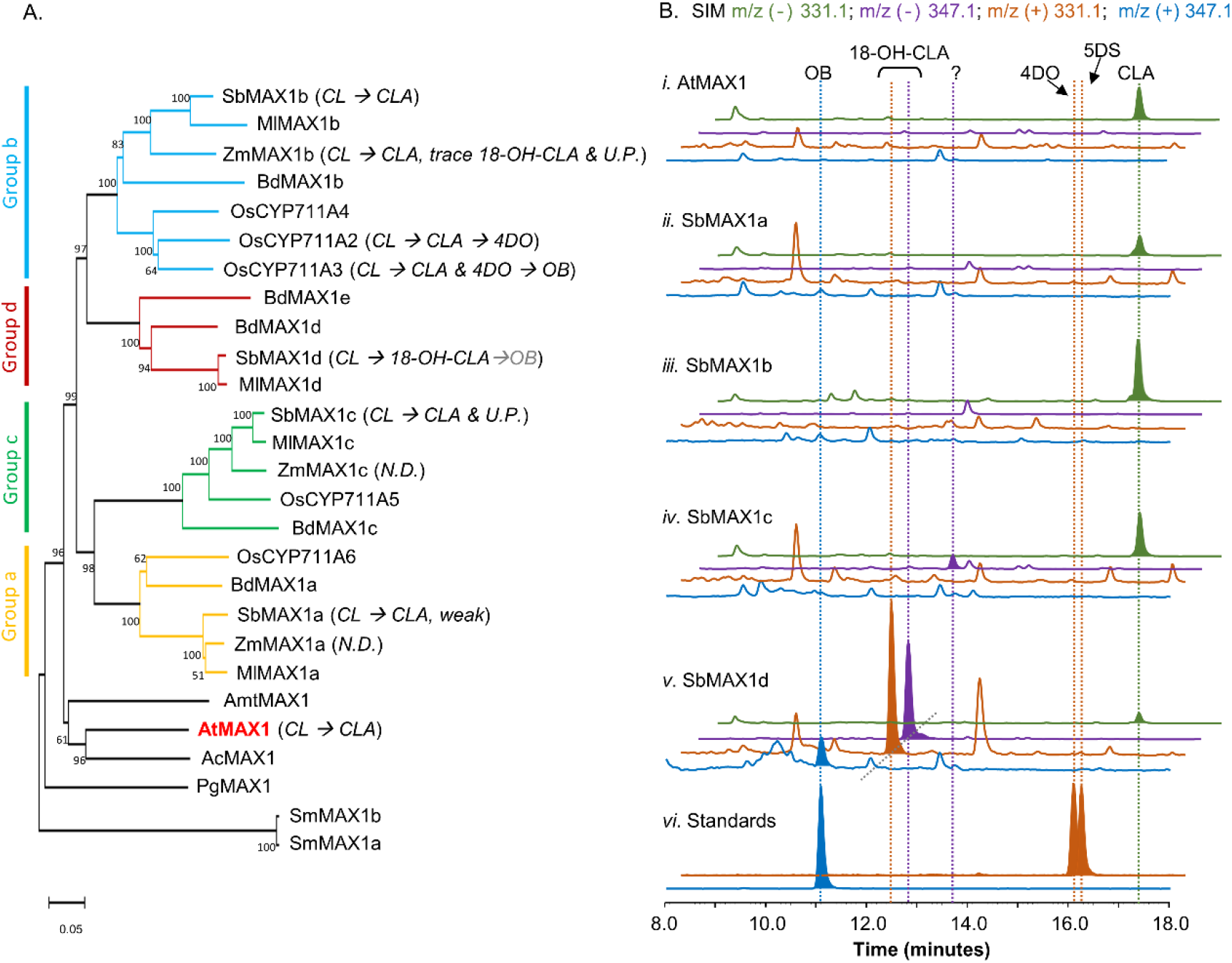
Functional characterization of MAX1 analogs from *S. bicolor*. (a) Phylogenetic analysis of MAX1 analogs. The Phylogenetic tree was reconstructed in MEGA X using the neighbor-joining method based on amino acid sequence. The MAX1 analogs are from dicotyledons and monocotyledons. Species abbreviations: Sb, *Sorghum biocolor*; Ml, *Miscanthus lutarioriparius*; Zm, *Zea mays*; Bd, *Brachypodium distachyon*; Os, *Oryza sativa*; Amt, *Amborella trichopoda*; At, *Arabidopsis thaliana*; Ac, *Aquilegia coerulea*; Pg, *Picea glauca*; Sm, *Selaginella moellendorffii*. For the accession numbers of proteins, see Table S4. (b) SIM extracted ion chromatogram (EIC) at m/z^−^=331.1 (green), 347.1 (purple), and m/z^+^=331.1 (orange), 347.1 (blue) of CL-producing *E. coli* cocultured with *A. thaliana* P450 reductase 1 (ATR1)-expressing yeast i) expressing AtMAX1, ii) – v) expressing SbMAX1a-d, and vi) standards of OB, 4DO, and 5DS. CLA shows characteristic m/z^−^=331.1 (MW=332.40, [C_19_H_24_O_5_-H]^−^=[C_19_H_23_O_5_]^−^=331.1); 18-OH-CLA shows characteristic m/z^−^=347.1 and m/z^+^=331.1 (MW=348.40, [C_19_H_24_O_6_-H]^−^=[C_19_H_23_O_6_]^−^=347.1, [C_19_H_24_O_6_-H_2_O+H]^+^=[C_19_H_23_O_5_]^+^=331.1); OB shows characteristic m/z^+^=347.1 (MW=346.38, [C_19_H_22_O_6_+H]^+^=[C_19_H_23_O_6_]^+^=347.1); 4DO and 5DS show characteristic m/z^+^ signal (MW=330.38, [C_19_H_22_O_5_+H]^+^=[C_19_H_23_O_5_]^+^=331.1). All traces are representative of at least three biological replicates for each engineered *E. coli*-*S. cerevisiae* consortium. 18-OH-CLA stands for 18-hydroxy-CLA. MW stands for molecular weight. Strain used for analysis: AtMAX1 (**ECL/YSL1**, Table S2), SbMAX1a-d (**ECL/YSL2a-d**, Table S2).

Here, harnessing the recently developed SL-producing microbial consortia^19^ (Figure S2), we investigated SL biosynthesis in *Sorghum bicolor*, which turns out to be distinct from that in rice^15^. We identified SbMAX1d as a unique CYP that catalyzes up to four oxidation steps converting CL to 18-hydroxy-CLA and a small amount of OB. Following this discovery, we found the substrate of LGS1 is likely 18-hydroxy-CLA. The addition of the sulfo group to 18-hydroxy can inhibit further oxidation towards the synthesis of OB, and 18-sulphate-CLA synthesized from LGS1 can spontaneously form comparable amount of 4DO and 5DS with sulphate functioning as an easier leaving group than the original hydroxyl. Our study discovered a second synthetic route towards the synthesis of (*S*)-type SL which employs the unique SOT LGS1. However, the enzyme catalyzing the exclusive conversion of 18-sulphate-CLA to 5DS is still missing and requires further investigation into sorghum (Figure 1).

## Results and Discussion

Same as other *Poaceae* family members, sorghum does not encode CYPs that belong to CYP722C subfamily but encode four MAX1 analogs. To understand the evolutionary relationship of these MAX1 homologues, we conducted a phylogenetic analysis of selected MAX1 analogs from dicotyledons and monocotyledons (Figure 2a, Figure S1). Noticeable, the MAX1 analogs from grasses fall into four different subclades, which are named group a-d here for simplicity (Figure 2a). Sorghum’s four MAX1 analogs fall into each of the four groups, while maize and rice only encode MAX1 analogs from group a-c but not group d. To understand the biosynthetic machinery of 5DS and OB in sorghum, MAX1 analogs from sorghum (Table S1) were introduced to the CL-producing microbial consortia (**ECL**, Table S2, Figure 2b). Interestingly, expression of SbMAX1d to CL-producing consortium (**ECL/YSL2d**, Table S2) led to the synthesis of OB and 18-hydroxy-CLA (verified through high-resolution mass spectrometry [HRMS] analysis, Figure S3a). The production of 18-hydroxy-CLA by SbMAX1d is much more efficient than all the SL synthetic CYPs we examined previously (CYP722Cs and OsCYP711A2, resulting in **ECL/YSL3-5** Table S2, Figure 2b, Figure S4)^18^. Likely SbMAX1d first catalyzes three-step oxidation on C19 to synthesize CLA, followed by two-step oxidation on C18 to afford the synthesis of 18-hydroxy-CLA and subsequently 18-oxo-CLA, which can spontaneously convert to OB (Figure 1)^18, 20^. Whether the conversion from 18-hydroxy-CLA to OB is catalyzed by SbMAX1d as shunt product, or by endogenous enzymes in yeast or *E. coli* remains to be investigated.

In addition, SbMAX1c converted CL to CLA and one new peak of molecular weight same as 18-hydroxy-CLA (16Da more than that of CLA) (Figure 2b, S3b). However, due to the low titer of SLs from the microbial consortia and the lack of commercially available standards, we cannot verify the identities of this compound synthesized by SbMAX1c currently. The other two MAX1 analogs examined simply catalyze the conversion of CL to CLA without further structural modifications (Figure 2b). The MAX1 analogs were also introduced to **ECL/YSL2d** or **ECL/YSL5** that produce 18-hydroxy-CLA and OB or 5DS (resulting strain: **ECL/YSL6-7**, Table S2) but no new conversions were detected (Figure S5). The newly discovered and unique activities of SbMAX1d and SbMAX1c imply the functional diversity of MAX1 analogs encoded by monocot plants, with much remains to be investigated.

To synthesize 5DS by Group II CYP722C (or 4DO by OsCYP711A2), likely C19 functions as the nucleophile to attack C18, which enables C18-hydroxy to recruit one proton and form water as the leaving group (Figure S6)^15, 17^. However, hydroxy group is generally not a favorable leaving group, and it often needs to be activated to trigger the subsequent reactions (e.g., intramolecular cyclization). Common hydroxy activation strategies used in nature include acetylation, phosphorylation, and sulfonation^21–23^. Sulfation/intramolecular cyclization has been reported to be employed in microbial natural product biosynthesis, such as ficellomycin from *Streptomyces ficellus*^23^, but seldom in plant. The discovery of the unique SbMAX1d synthesizing 18-hydroxy-CLA as the major product leads to the hypothesis that LGS1 may modify the 18-hydroxyl group to form 18-sulphate-CLA, which will prohibit further oxidation towards the formation of OB and promote the nucleophilic attack on C18 to form C ring. Introduction of LGS1 to **ECL/YSL2d** (resulting **ECL/YSL8a**, Table S2) resulted in substantial decrease of 18-hydroxy-CLA and the appearance 4DO and 5DS (ratio ~ 1:1, Figure 3a), though the amount is low in comparison to 18-hydroxy-CLA and OB (Figure 3a). Similar to many previous sulfotransferase studies^24^, 18-sulphate-CLA was not detected from *in vivo* assays using SL-producing microbial consortia (Figure S7). 4DO and 5DS are synthesized in similar levels, which indicates that the conversion from 18-sulphate-CLA to the canonical SL structures are likely spontaneous with 18-sulphate as an easier leaving group than water formed from 18-hydroxy (Figure S8). There is likely other enzyme(s) involved downstream of or simultaneous with LGS1 to guarantee the conversion of 18-sulphate-CLA to 5DS exclusively instead of a 4DO/5DS mixture. We thus examined the function of SbMAX1a, 1b, 1c, SbCYP722B, SbCYP728B35, SbCYP728B1, and ZmCYP728B35 in the 4DO/5DS/18-hydroxy-CLA-producing consortium **ECL/YSL8a** (resulting **ECL/YSL9-10**, Table S2)^25^. Unfortunately, we were unable to see any changes to the ratio between 5DS and 4DO (Figure S9). Further genomics-based analysis on Sorghum is required to identify the missing components that is responsible for the inversion of the stereochemistry on the C ring.

**Figure 3.**
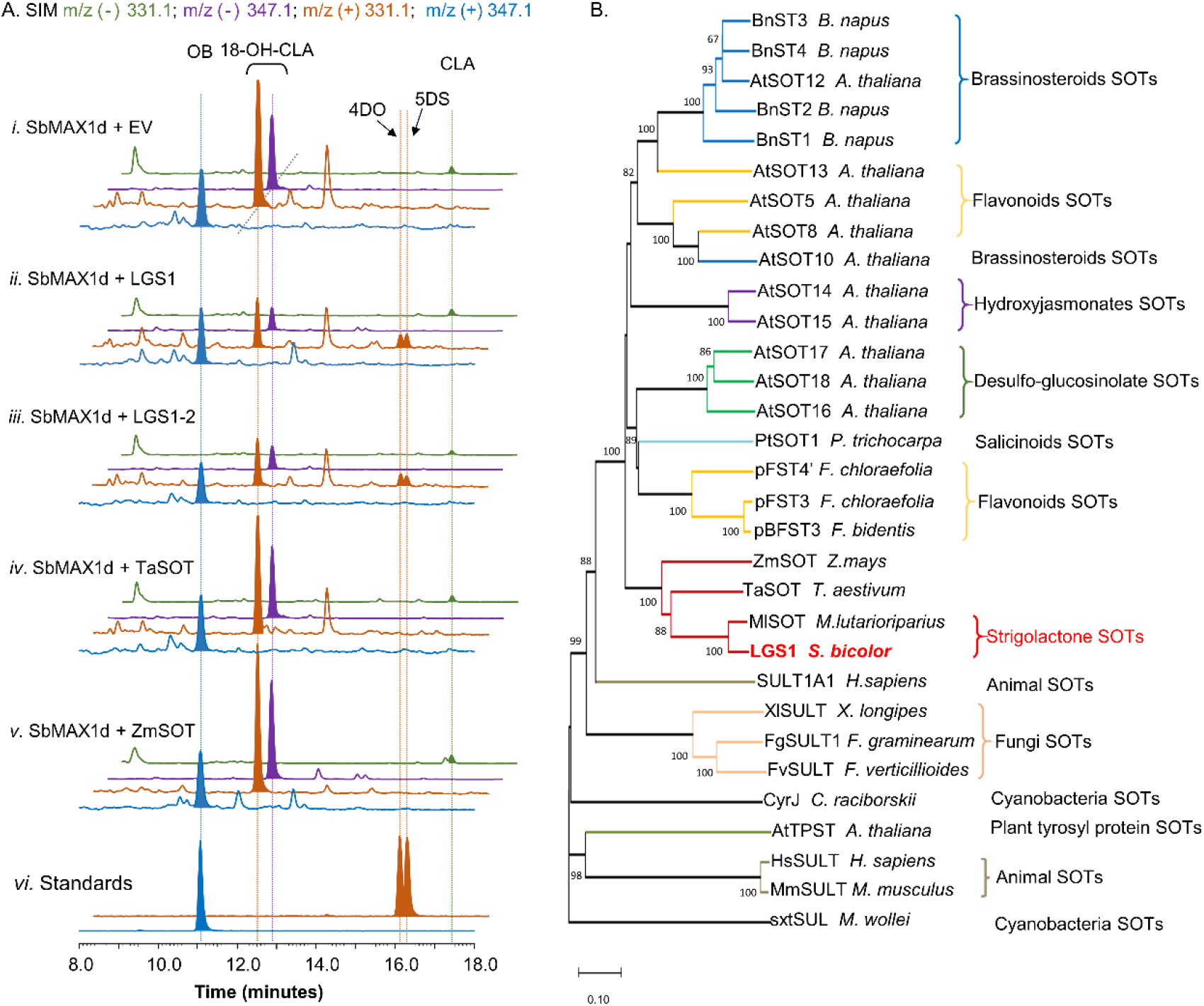
Functional characterization of LGS1 and analogs using CL-producing microbial consortium expressing SbMAX1d. (a) SIM EIC at m/z^−^=331.1 (green), 347.1 (purple), and m/z^+^=331.1 (orange), 347.1 (blue) of CL-producing *E. coli* cocultured with yeast expressing ATR1, SbMAX1d and i) empty vector (EV), ii) LGS1, iii) LGS1-2, iv) SOT from *Triticum aestivum* (TaSOT), v) SOT from *Zea mays* (ZmSOT), and vi) standards of OB, 4DO and 5DS. All traces are representative of at least three biological replicates for each engineered *E. coli*-*S. cerevisiae* consortium. (b) Phylogenetic analysis of LGS1. The Phylogenetic tree was reconstructed in MEGA X using the neighbor-joining method based on amino acid sequence. The SOTs are from animals, plants, fungi and cyanobacteria. For the accession numbers of proteins, see Table S5. The sequence of LGS1 is from sorghum WT Shanqui Red, LGS1-2 variation is a reference sequence from NCBI, and is four amino acids (DADD) longer than LGS1, see Table S6.

To further validate the proposed mechanism of LGS1 in Sorghum SL biosynthesis (Figure S8), lysates from yeast expressing LGS1 were incubated with spent medium of CL-producing consortia expressing SbMAX1d. When LGS1 was assayed with 18-hydroxy-CLA and PAPS, 18-hydroxy-CLA was nearly completely consumed. 4DO and 5DS were observed, but not 18-sulphate-CLA, which is likely due to the low stability (Figure 4). The addition of PAPS to the lysate assay system results in enhanced consumption of 18-hydrxoy-CLA and also synthesis in 4DO/5DS (Figure 4), which indicates that LGS1 is a PAPS-dependent sulfotransferase.

**Figure 4.**
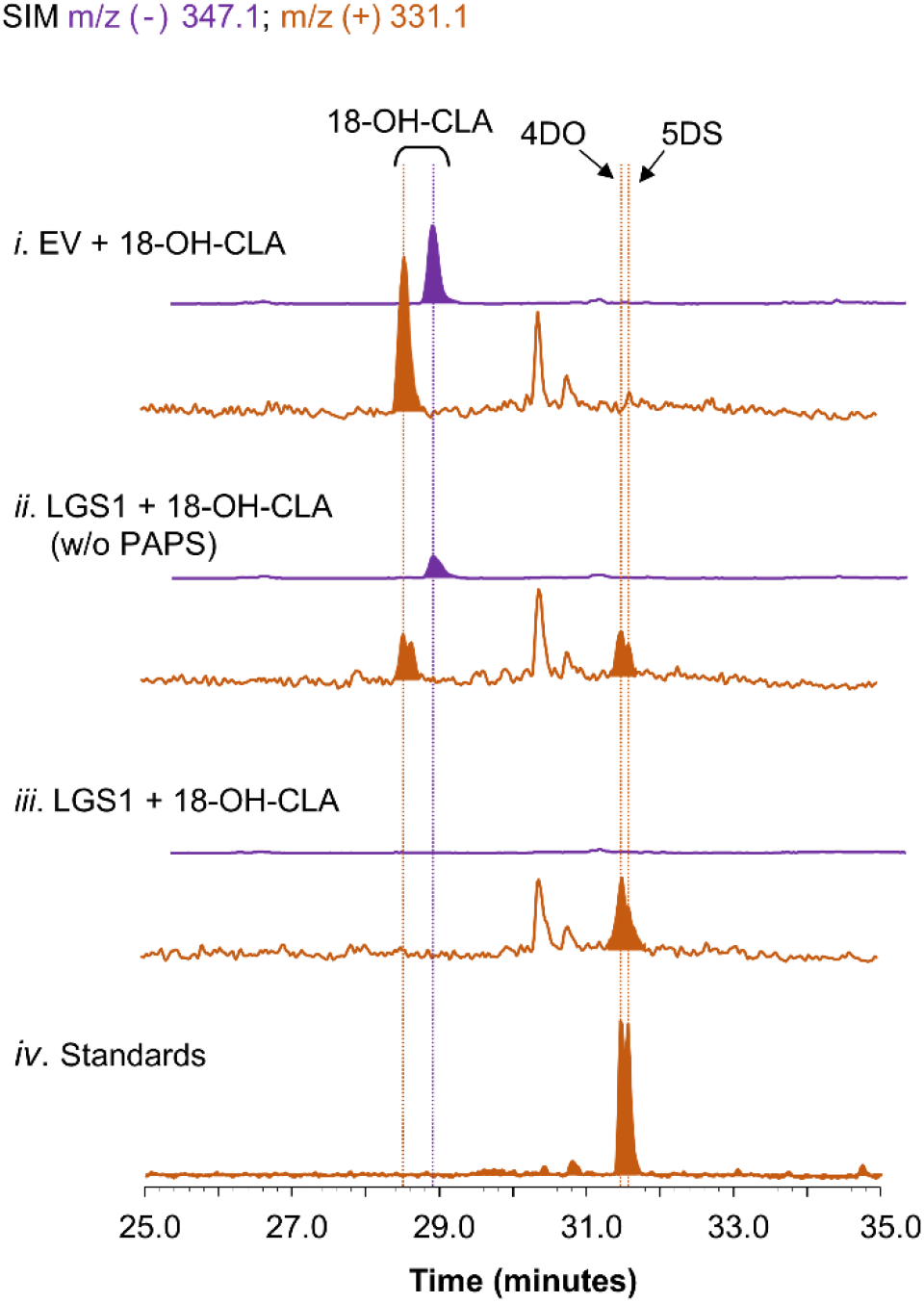
Characterization of LGS1 activity using crude lysate assay. SIM EIC at m/z^−^=347.1 (purple) and m/z^+^=331.1 (orange) of crude lysate assay using i) EV-harboring yeast with PAPS, ii) LGS1-expressing yeast without PAPS, iii) LGS1-expressing yeast and PAPS, iv) authentic standard of 4DO and 5DS. The reaction was incubated for 1 hour with extracts of **ECL/YSL2d** medium, the samples were analyzed using Separation Method II (extraction method see Material and Methods in Supporting Information).

Similar to other plant SOTs, LGS1 is predicted to be localized in cytoplasm. Cytosolic SOTs contain several conserved PAPS-binding motifs, including the one interacts with 5′-phosphate of PAPS (TYPKSGT), 3’-phosphate of PAPS (YxxRNxxDxxVS), and nucleotide of PAPS (GxxGxxK/R)^26^. Multiple sequence alignment indicates that LGS1 contains these motifs, but with some variations (<u>SL</u>PKSGT and YxxR<u>E</u>xxDxxVS, respectively) (Figure S10). LGS1 contains the highly conserved histidine residues (H216)^27^ and moderately conserved histidine residues (H317A) (Figure S10) which likely act as a base to remove the proton from the substrate hydroxyl group, thereby forming an oxygen anion, and then attacking the sulfo group of PAPS to complete the transfer of the sulfo group. To determine whether these residues play a key role in catalysis, we conducted site directed mutagenesis on residues likely act as a catalytic base (H216A, H317A) or crucial for PAPS binding (K148A, Y247F)^26^. While LGS1^H216A^ (resulting strain: **YSL8f**, Table S2) exhibited same activity as wildtype LGS1, replacing LGS1 with LGS1^K148A^, LGS1^Y247F^, and LGS1^H317A^ in **ECL/YSL8a** (resulting strain: **YSL8g-i**, Table S2) completely abolished the synthesis of 4DO and 5DS (Figure S11), implying that these residues are critical to the catalytic activity of LGS1 (Figure S11).

SOTs universally exist in all types of organisms and involve in the modification of both small molecules (e.g., steroids^28^) and macromolecules (e.g., glycosaminoglycans^29^). Among various plant SOTs, the ones from *Arabidopsis thaliana* are the most studied, with 10 out of 21 AtSOTs of known functions or substrates^24, 30^. To examine if similar LGS1-involved SL biosynthetic mechanism exists in other plants, likely *Poaceae* plants, we used LGS1 protein sequence as a query to seek for LGS1 analogs (Table S5). We were only able to find one sulfotransferase from *Miscanthus lutarioriparius* (MlSOT, 401 a.a., 80% identity) of high similarity to LGS1 (452 a.a.), while the next few on the list are all quite different from LGS1. We selected a few SOTs that exhibit highest similarity to LGS1 including MlSOT, SOTs from *Triticum aestivum* (TaSOT, 345 a.a., 55% identity), and *Zea mays* (ZmSOT, 451 a.a., 53% identity), and tested the activity in **ECL/YSL8c-e** (Table S2). As expected, only MlSOT was able to synthesize 5DS and 4DO but with a much lower efficiency than LGS1 (Figure S11), while ZmSOT and TaSOT didn’t change the SL production profile (Figure 3a). To further understand the evolutionary relationship between LGS1 and other plant SOTs, we constructed a phylogenetic analysis of various SOTs from plants, animals, bacteria, and fungal (Table S5, Figure 3b). As expected, LGS1 belong to plant SOT family, but are distinct from other characterized plant SOTs^24^. LGS1 and MlSOT are located on a unique subbranch that is different from all the other plant SOTs (Figure 3b).

Multiple independent natural LGS1 loss-of-function varieties have been found in *Striga*-prevalent areas in Africa and are rare outside of *Striga*-prone region, which indicates that the lack of *LGS1* gene can adapt to weed parasitism^31^. *M. lutarioriparius* encode four MAX1 analogs, and each exhibits high similarity and corresponds to one of the four SbMAX1s^32^. Because MlSOT also exhibits the same activity as LGS1, highly likely *M. lutarioriparius* harness the same LGS1-involving strategy and produces similar SL profiles to sorghum. However, the lack of LGS1 paralogs in other crops (e.g., maize) implies that much remains to be characterized about SL biosynthesis in these economically significant plants. For example, maize has been reported to produce 5DS and non-classical SLs but not (*O*)-type SLs^33–35^. However, same as other members from the *Poaceae* family, maize does not encode CYP722C analogs. The lack of LGS1 functional paralog thus indicates that a different synthetic route towards 5DS remains to be uncovered from maize. The activities of MAX1 analogs from maize (Table S1 and S3) were examined in different microbial consortia, as well (**ECL/YSL11**, Table S2). ZmMAX1b^36^ exhibited similar activity to SbMAX1c: in addition to converting CL to CLA, it produced trace amounts of 18-hydroxy-CLA and an unknown oxidated product as SbMAX1c (Figure S12). ZmMAX1a and c showed no activity towards CL (Figure S12). Our results suggest that the 5DS biosynthesis in maize likely requires unknown types of enzymes yet to be identified.

## Conclusion

Taken together, we clarified the function of LGS1, revealed the cryptic sulfonation in the formation of the canonical SLs skeleton, and explained how the participation of LGS1 redirects the biosynthetic pathway of canonical SLs, thereby switch the product from OB into 5DS.

## Materials and Methods

### Reagents and general procedures

(±)5-deoxy-strigol (purity >98%) and (±)-orobanchol were purchased from Strigolab (Italy), (±)4-deoxyorobanchol (also named as (±)-2’-epi-5-deoxystrigol) were purchased from Chempep Inc. (USA). Adenosine 3′-phosphate 5′-phosphosulfate lithium salt hydrate (PAPS) and the antibiotics were purchased from Sigma-Aldrich Co. (USA). The BP Clonase II Enzyme Mix, LR Clonase II Enzyme Mix, and Gateway pDONR221 vector were obtained from Invitrogen (USA). The *S. cerevisiae* Advanced Gateway Destination Vector Kit were obtained from Addgene. Expand high fidelity PCR system (Roche Life Science) were used for PCR reactions (Bio-Rad,USA). The *Escherichia coli* TOP 10 competent cells were purchased from Life Technologies. The genes were synthesized by IDT (Coralville, IA) and primers were synthesized by Life Technologies (USA). DNA sequencing were performed at Genewiz (San Diego, CA).

For CL production, XY medium [13.3 g/L KH_2_PO_4_, 4 g/L (NH_4_)_2_HPO_4_, 1.7 g/L citric acid, 0.0025 g/L CoCl_2_, 0.015 g/L MnCl_2_, 0.0015 g/L CuCl_2_, 0.003 g/L H_3_BO_3_, 0.0025 g/L Na_2_MoO_4_, 0.008 g/L Zn(CH_3_COO)_2_, 0.06 g/L Fe(III) citrate, 0.0045 g/L thiamine, 1.3 g/L MgSO_4_, 5 g/L yeast extract and 40 g/L xylose, pH 7.0] was prepped and used as previously described^19^. For yeast ectopic expression, synthetic dropout (SD) medium was used [0.425g yeast nitrogen base (YNB, BD Biosciences, USA), 1.25g (NH_4_)_2_SO_4_ dissolved in 200 mL distilled water (dH_2_O), autoclave at 121 °C for 20 min. Add 25 mL 200 g/L glucose and 25 mL 20 g/L amino acid drop-out mix (Clontech. Inc., Japan) solution to prepare the medium]. LC-MS was carried out on a Shimadzu LC-MS 2020 (Kyoto, Japan) with LC-MS grade solvent. High resolution mass spectrometry (HR-MS) analysis was carried on a Synapt G2-Si quadrupole time-of-flight mass spectrometer (Waters) coupled to an I-class UPLC system (Waters).

### Plasmid construction

All genes were codon optimized for *S. cerevisiae* (Table S6), synthesized, and cloned into the entry vector pDONR221 (Invitrogen) through Gateway® BP reaction. The genes were then introduced to the yeast expression vector through Gateway® LR reaction using destination vectors from Yeast Gateway Kit^37^. LGS1 mutants were constructed through PCR using primers listed in Table S7. PCR was performed using pAG416GPD-LGS1 as the template with Expand high fidelity PCR system. The amplified DNA fragment was purified, recovered using Gibson assembly, and used to transform TOP 10 competent cells. The sequence was verified using DNA sequencing.

### Culture conditions for *E. coli*-yeast consortium-based SL production

The *E. coli* strain ECL for carlactone production (Table S2) were prepared as described previously^19^. Single colony was grown overnight at 37 °C in 1 mL Luria-Bertani (LB) containing 25 μg/mL chloramphenicol, 50 μg/mL spectinomycin and 100 μg/mL ampicillin. 500 μL of the overnight culture was then used to inoculate 5 mL of fresh LB with the corresponding antibiotics and cultured at 37 °C and 220 rpm in the 100 mL Erlenmeyer flask. When OD600 reached ~0.6, IPTG was added with the final concentration at 0.2 mM, with ferrous sulfate supplemented at the same time (final concentration at 10 mg/L). Then the cultures were incubated at 22°C and 220 rpm for 15 hours. At the same time, single colony of each yeast strain harboring the corresponding cytochrome P450-expression constructs was used to inoculate 1 mL synthetic dropout medium (SDM). The seed culture was incubated at 28 °C and 220 rpm overnight. 100 μL of the overnight grown seed culture was used to inoculate 5 mL of the corresponding SD medium in a 100 mL Erlenmeyer flask and grown at 28 °C for 15 hours. The *E. coli* and yeast cells were harvested by centrifugation at 3,500 rpm for 5 min. Then they were re-suspended together in 5 mL of XY media and grown at 22°C for 60 hours.

### Metabolite extraction and analysis

Collect the cell pellet through centrifugation in a 50 mL centrifuge tube at 4,000 rpm for 10 min. The supernatant is transferred to another 50 mL centrifuge tube. The cell pellets were resuspended in 150 μL dimethylformamide (DMF) and 850 μL acetone, vortexed for 15 min, and centrifuged at 12,000 rpm for 10 min. The medium was extracted with 4 mL of ethyl acetate (EtOAc), vortex for 1 min, and centrifuged at 4,000 rpm for 20 min. The supernatant from both portions were collected and transferred to 1.7 mL microcentrifuge tubes, dried in vacufuge under reduced pressure (Eppendorf concentrator plus), and re-dissolve in 100 μL acetone. The extract was centrifuged at 12,000 rpm for 10 min before *LC-MS* analysis. The *LC-MS* analysis were performed on a C18 column (Kinetex® C18, 100 mm × 2.1 mm, 100Å, particle size 2.6 μm; Phenomex, Torrance, CA, USA). Separation Method I, the parameters were set as follows: column temperature: 40 °C, flow rate: 0.4 mL/min; mobile phase A: water containing 0.1% (v/v) formic acid; mobile phase B: acetonitrile containing 0.1% (v/v) formic acid. The LC program was set as follows: 0-28 min, 5%-100% B; 28-35 min, 100% B; 35-40 min, 5% B.

### Heterologous expression of recombinant LGS1 in *S. cerevisiae*

We first expressed LGS1 from *E. coli* strain BL21 (DE3) but failed. Then LGS1 was expressed from *S. cerevisiae* CEN.PK2 using low-copy number plasmid (pAG416GPD-LGS1, Table S1). One fresh colony of the LGS1-expressing yeast strain was first cultured in 1mL SDM lacking uracil (SD-Ura) medium, grown at 30°C and 220 rpm for overnight in a shaker incubator. 80 μL of the overnight culture was used to inoculate 5 mL SD-Ura medium (OD_600_ ≈ 0.1), grown at 30°C and 220 rpm for 48 h (OD_600_ ≈ 20). Cell pellets were then harvested by centrifuging at 3,500 rpm for 2 min, washed with 1 mL of water, and resuspend in 120 μL of 20 mM sodium phosphate buffer (pH=7.4). 50 μL of silicon beads [0.5 mm, Research Products International (RPI)] were then added to the cell suspension, which is then chilled on ice, and lysed using cell disruptor (FastPrep^®^-24, MP Biomedicals), The parameters were set as speed: 4.0 m/s, time: 30 s. The homogenate was centrifuge at 13,000 rpm for 2 min, and the supernatant was used for the crude lysate-based enzyme assays.

### Yeast crude lysate-based enzyme assays

50 μL of crude enzyme extract mentioned above is incubated with 5 μL of concentrated metabolic extract dissolved in DMF (extracted from 3 mL co-culture strain), with or without 100 μM PAPS, and incubated at 30°C for 1 hour. Enzyme assay using yeast strain expressing an empty vector as the negative control. The reaction mixture was quenched by adding an equal volume of acetonitrile followed by vigorous vortexing to remove the protein. The quenched reaction mixtures were then centrifuged at 13,000 rpm for 2 min. 17 μL of supernatant was subjected to LC*-*MS analysis with the C18 column (Kinetex® C_18_, 100 mm × 2.1 mm, 100Å, particle size 2.6 μm; Phenomex, Torrance, CA, USA). In order to detect speculative 18-sulphate-CLA, an intermediate with an increased polarity, we use a different separation method: Separation Method II. The parameters were set as follows: column temperature: 40 °C, flow rate: 0.4 mL/min; mobile phase A: water containing 0.1% (v/v) formic acid; mobile phase B: acetonitrile containing 0.1% (v/v) formic acid. The LC program was set as follows: 0-3 min, 5%-11% B; 3-13 min, 11%-19% B; 13-21 min, 19%-27.5% B; 21-24 min, 27.5%-34% B; 24-28 min, 34%-42% B; 28-32min, 42%-90% B; 32-34 min, 90%-100% B; 34-35.5 min, 100%-5% B; 35.5-40 min, 5% B.

## ASSOCIATED CONTENT

### Supporting Information

Supplementary figures on sorghum SL biosynthesis characterization and LGS1 identification, supplementary tables summarizing plasmids, strains, and gene sequences used in this study (PDF)

## AUTHOR INFORMATION

### Author Contributions

S.W and Y.L. conceived the project; S.W. and Y.L. designed the experiments; S.W. performed the experiments and analyzed the results; S.W and Y.L. wrote the manuscript.

## ACKNOWLEDGMENT

We thank the Metabolomics Core Facility at UC Riverside and Dr. Anil Bhatia for instrument access, training, and data analysis; S. Xu for studying protein-protein interaction of SL biosynthetic enzymes identified in this study; A. Zhou for the construction of SYL89; and Dr. K. Zhou for the valuable feedback in the preparation of the manuscript.

## ABBREVIATIONS

5DS: 5-deoxystrigol
4DO: 4-deoxyorobanchol
OB: orobanchol
SL: Strigolactone
CL: carlactone
CLA: carlactonoic acid
SOT or SULT: sulfotransferase
PAPS: 3′-phosphoadenosine 5′-phosphosulfate
CYP: cytochrome P450
MAX1: MORE AXILLARY GROWTH 1
LGS1: LOW GERMINATION STIMULANT 1
HRMS: high-resolution mass spectrometry

