## Supplementary material for "A unique sulfotransferase-involving strigolactone biosynthetic route in Sorghum": SI

Sheng Wu[1], Yanran Li[1*]

[1] Department of Chemical and Environmental Engineering, University of California, Riverside, California 92521, USA

**Correspondence should be addressed to Yanran Li*

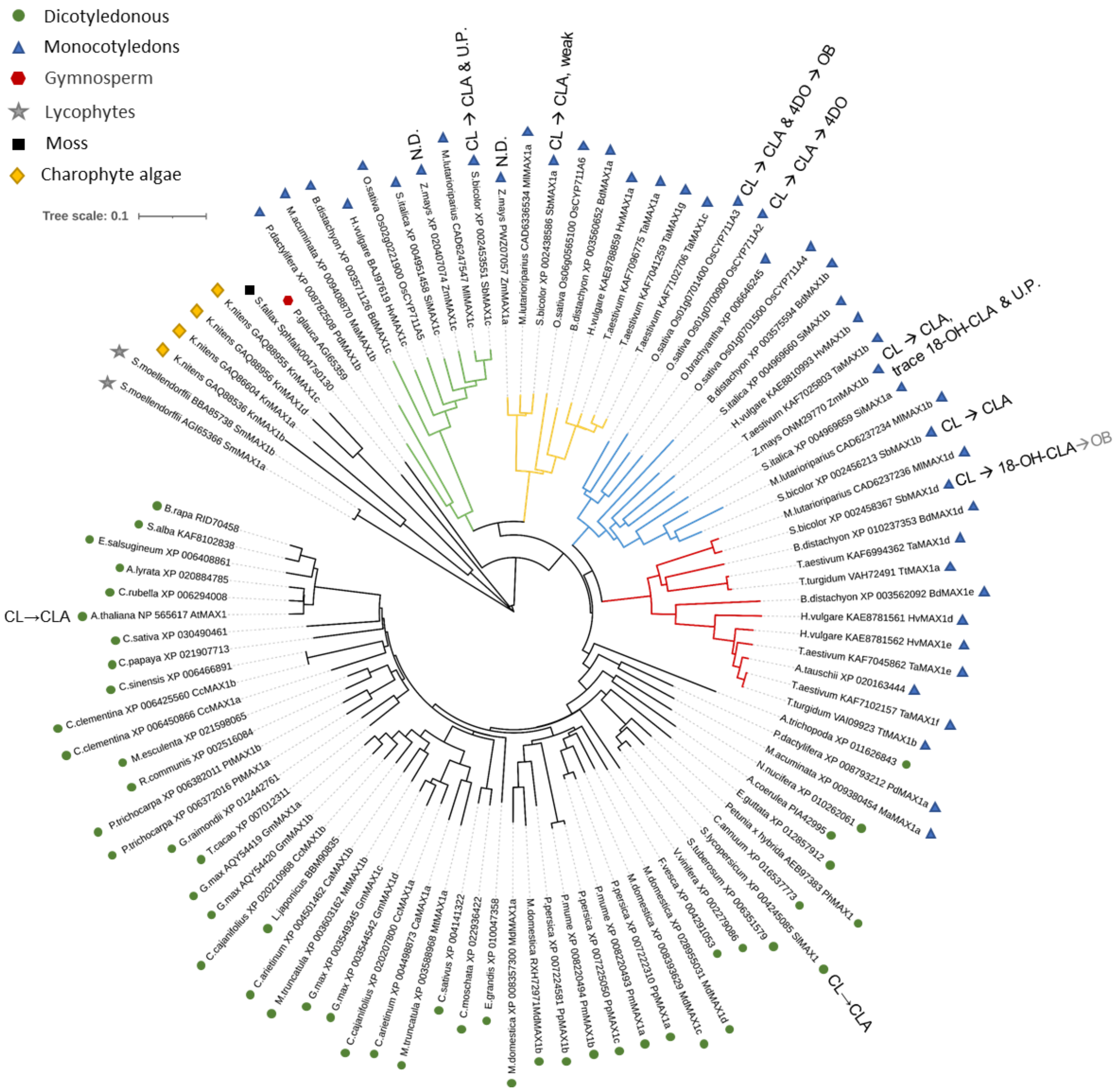


**Figure. S1.** Phylogenetic tree of MAX1 homologs. The phylogenetic tree was constructed by MEGA X program with neighbor joining method based on amino acid sequence (80% partial deletion, 1000 bootstraps, p-distance mode). This analysis involved 102 CYP genes (Table S4) that belong to CYP711A clan. CYP711A from monocotyledons can be divided into four groups, which are highlighted in green, yellow, blue, and red, respectively. MAX1 analogs of confirmed functions were annotated, which can also be found in Table S3.

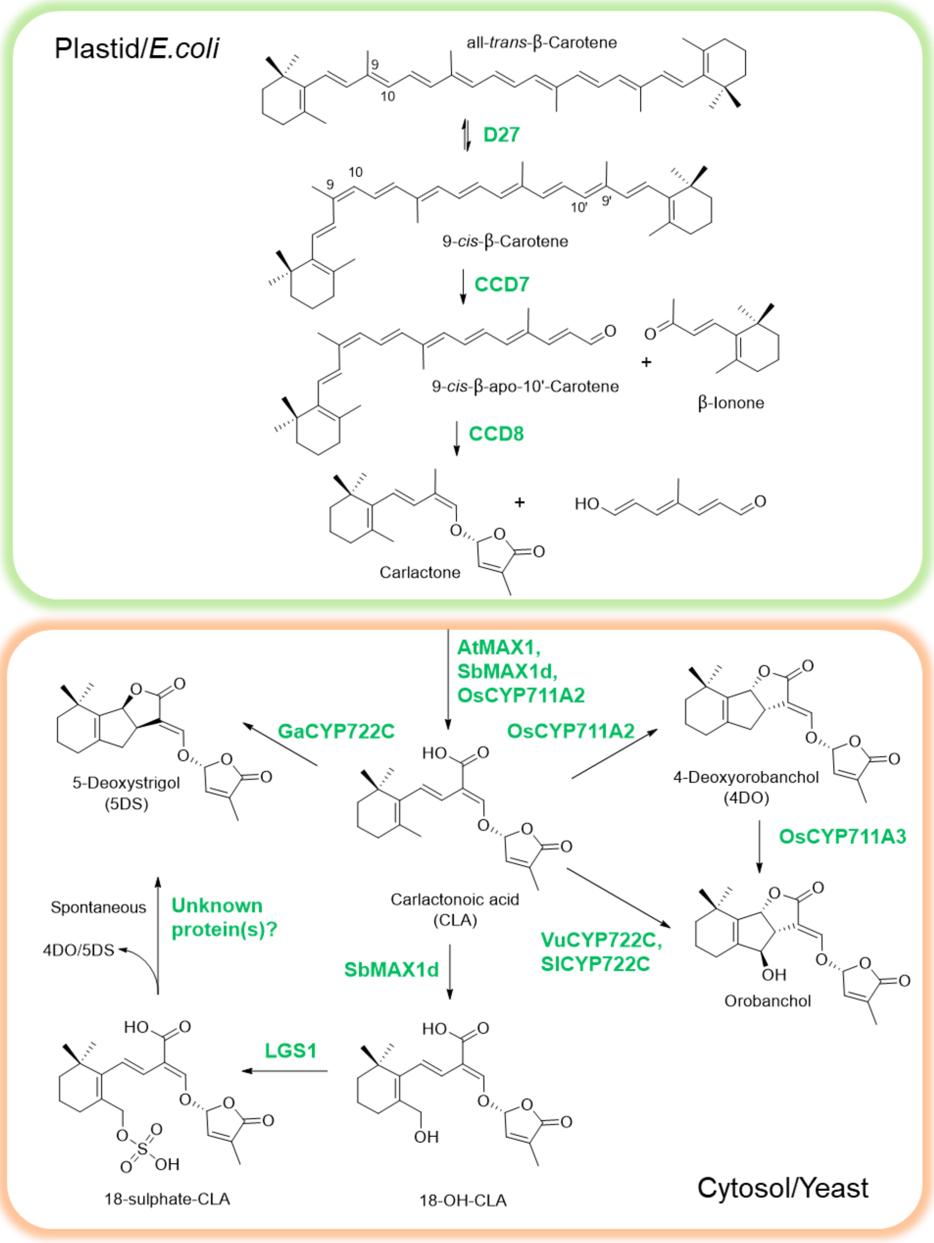


**Figure S2**. Illustration of SL-producing consortium. D27, DWARF27, a [2Fe-2S]-containing isomerase; CCD7, carotenoid cleavage dioxygenase 7; CCD8, carotenoid cleavage dioxygenase 8; MAX1, MORE AXILLARY GROWTH 1, belong to CYP711A1 subfamily; CYP711A2, cytochrome P450 711A2 subfamily; CYP711A3, cytochrome P450 711A3 subfamily; CYP722C, cytochrome P450 722C subfamily; At, *Arabidopsis thaliana*; Os, *Oryza sativa*; Ga, *Gossypium arboreum*; Vu, *Vigna unguiculata*; Sl, *Solanum lycopersicum*; Sb, *Sorghum bicolor*.

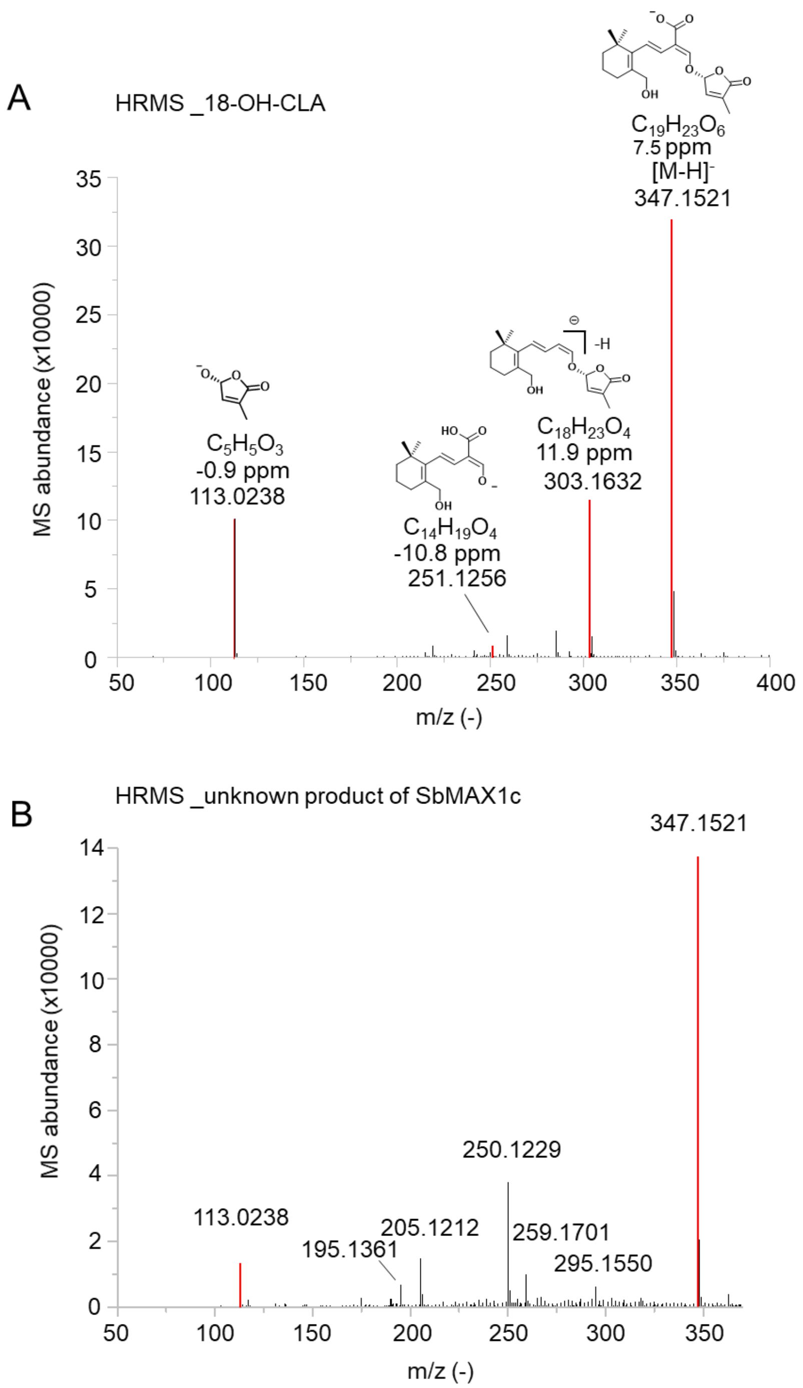


**Figure S3.** HRMS analysis of 18-hydroxy-CLA and the unknown product synthesized by SbMAX1c in *E. coli*-*S. cerevisiae* co-culture. (A) HRMS of 18-hydroxy-CLA produced by CL-producing consortia **ECL/YSL2d** (Table S2). The main fragment ions at m/z 113, 251 and 303 was consistent with the data in literature[1]. (B) HRMS of the unknown compound produced by CL-producing consortia **ECL/YSL2c** (Table S2).

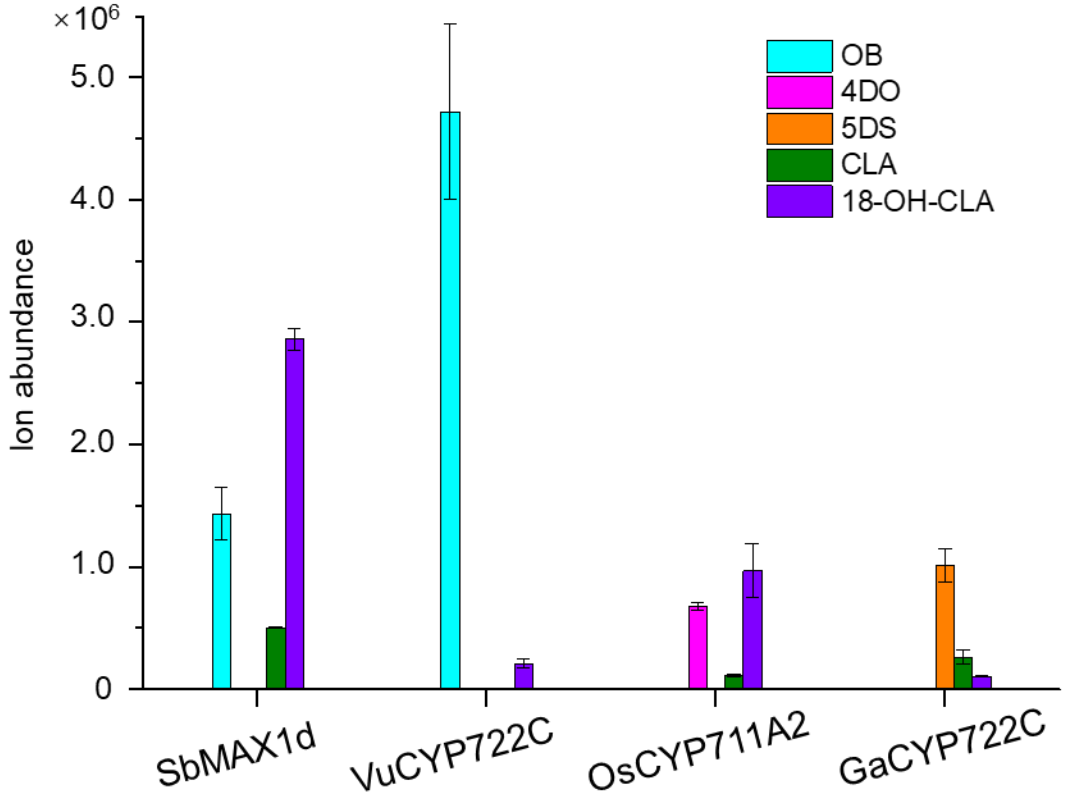


**Figure S4.** Comparison of 18-hydroxy-CLA and OB synthesis by SbCYP711A1, OsCYP711A2, VuCYP722C, and GaCYP722C in microbial consortia **ECL/YSL2d**, **ECL/YSL3**, **ECL/YSL4**, **ECL/YSL5**, respectively (Table S2). The error bars represent the s.d. of the replicates.

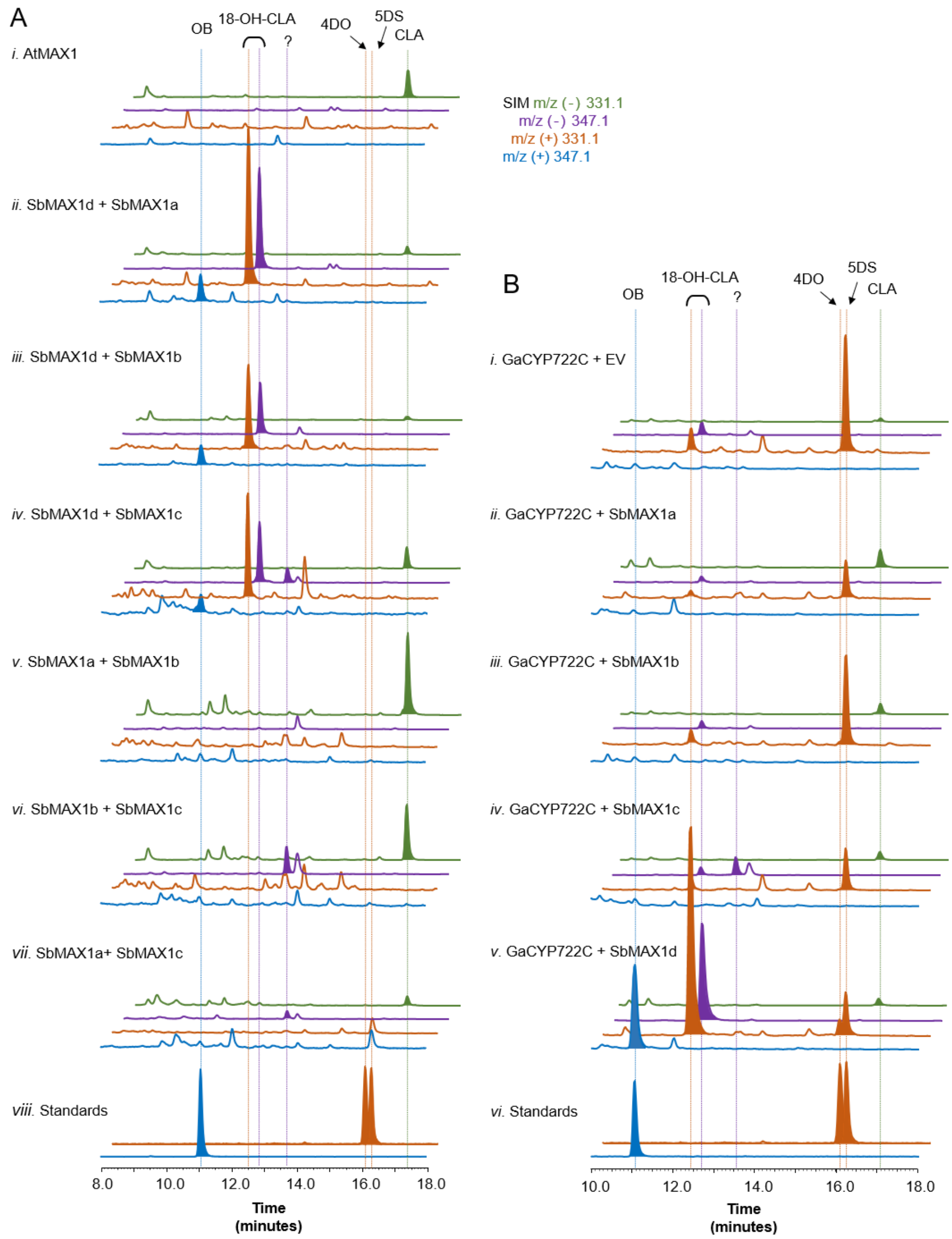


**Figure S5.** LC-MS analysis of metabolites produced by consortia expressing SbMAX1a-d. (A) SIM EIC at $m/z^-$=331.1 (green), 347.1 (purple), and $m/z^+$=331.1 (orange), 347.1 (blue) of **ECL** (Table S2) cocultured with ATR1-expressing yeast expressing i) AtMAX1 **(ECL/YSL1**, Table S2); ii) SbMAX1d and SbMAX1a; iii) SbMAX1d and SbMAX1b; iv) SbMAX1d and SbMAX1c; v) SbMAX1a and SbMAX1b; vi) SbMAX1b and SbMAX1c; vii) SbMAX1a and SbMAX1c (**ECL/YSL6a-f**, Table S2), and viii) standards of OB, 4DO and 5DS. (B) SIM EIC at $m/z^-$=331.1 (green), 347.1 (purple), and $m/z^+$=331.1 (orange), 347.1 (blue) of **ECL** cocultured with yeast expressing ATR1, AtMAX1, GaCYP722C and i) empty vector (EV, **ECL/YSL7N**, Table S2)), ii) SbMAX1a; iii) SbMAX1b; iv) SbMAX1c; v) SbMAX1d (**ECL/YSL7a-d**, Table S2), and vi) standards of OB, 4DO and 5DS.

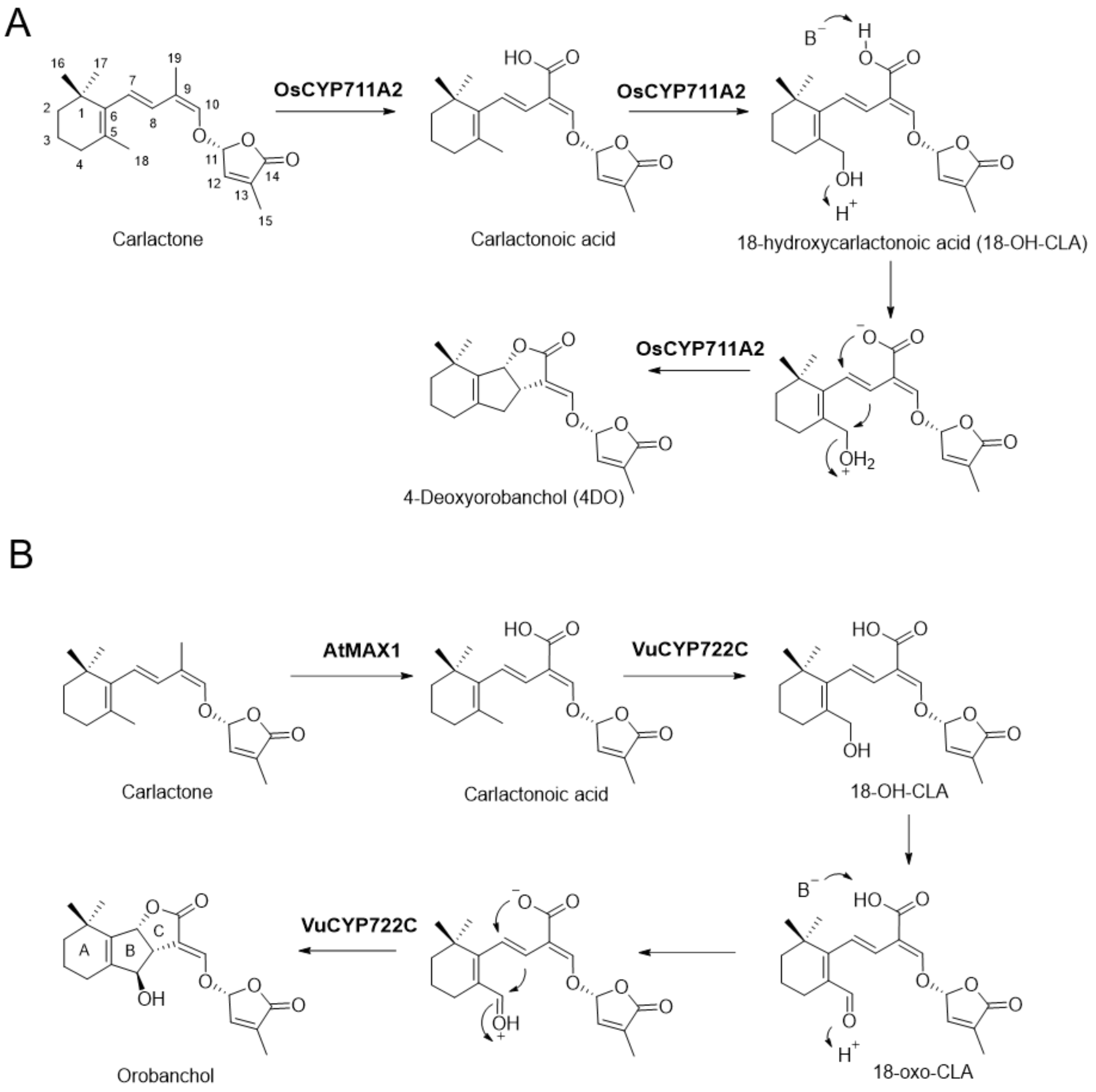


**Figure S6.** Proposed enzymic mechanism for the synthesis of 4DO, 5DS and OB by CYP711A and CYP722C. (A) The hydroxylation of CLA at C-18 position to form 18-hydroxy-CLA is essential for the formation of the BC ring. 18-hydroxy-CLA is likely the precursor of canonical SL, such as 4DO, 5DS and OB. To synthesize 4DO, a proton is likely transferred to the C-18 hydroxyl group, turning it into a leaving group, which then initiates the intramolecular nucleophilic substitution reaction accompanied by the elimination of water. The synthesis of 4DO and 5DS maybe very similar, except that the stereoselectivity of BC ring formation is different, and the formation of 5DS is catalyzed by GaCYP722C. (B) For OB biosynthesis, the 18-position hydroxyl group of CLA can be further oxidized to aldehyde (a reactive intermediate, 18-oxo-CLA) by VuCYP722C, which triggered the BC-ring closure. A nucleophilic addition-based cyclization strategy is used here.

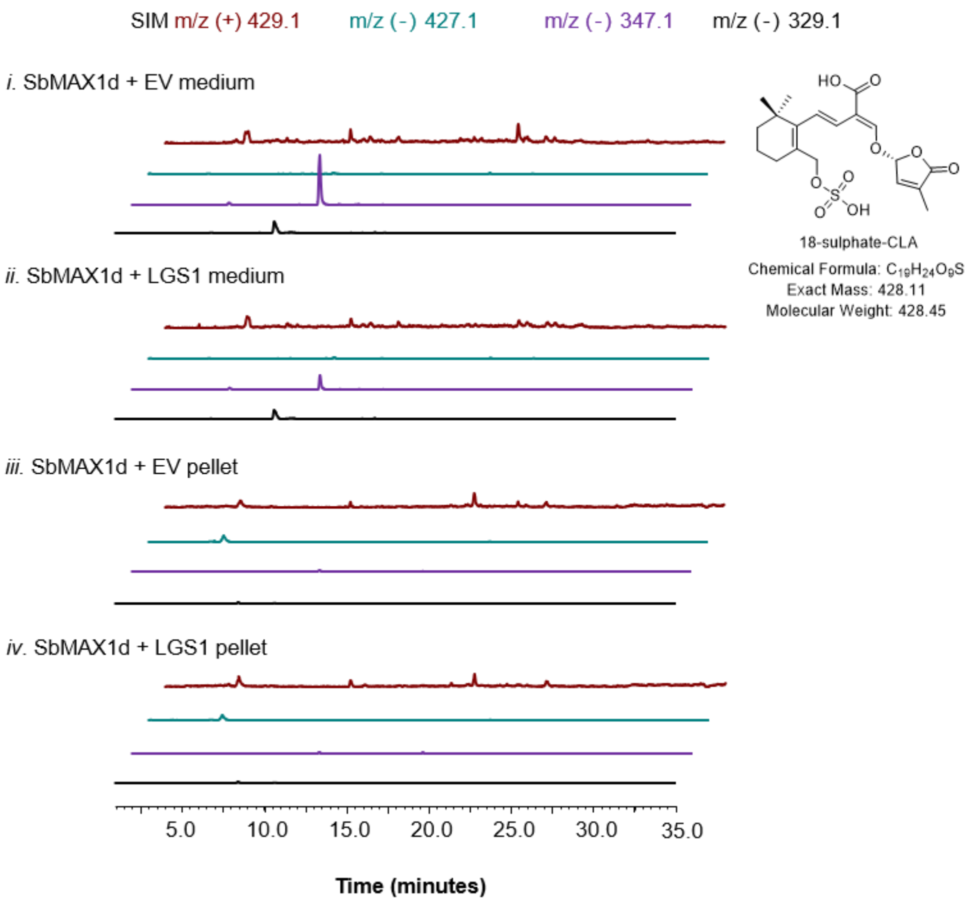


**Figure S7.** 18-sulphate-CLA was not detected from either the pellets or the medium of the microbial consortia. SIM-EIC using 18-sulphate-CLA's characteristic $m/z^+$ signal (MW=428.45, $[C_{19}H_{24}O_9S+H]^+=[C_{19}H_{25}O_9S]^+=429.1$), $m/z^-$ signal ($[C_{19}H_{24}O_9S-H]^-=[C_{19}H_{23}O_9S]^-=427.1$, $[C_{19}H_{24}O_9S-HSO_3]^-=[C_{19}H_{23}O_6]^-=347.1$, $[C_{19}H_{24}O_9S-H_3SO_4]^-=[C_{19}H_{21}O_5]^-=329.1$) of **ECL** (Table S2) cocultured with yeast expressing ATR1, SbMAX1d and i) EV (medium extracts, **ECL/YSL8N**, Table S2); ii) LGS1 (medium extracts, **ECL/YSL8a**, Table S2); iii) EV (cell pellet extracts); iv) LGS1 (cell pellet extracts)

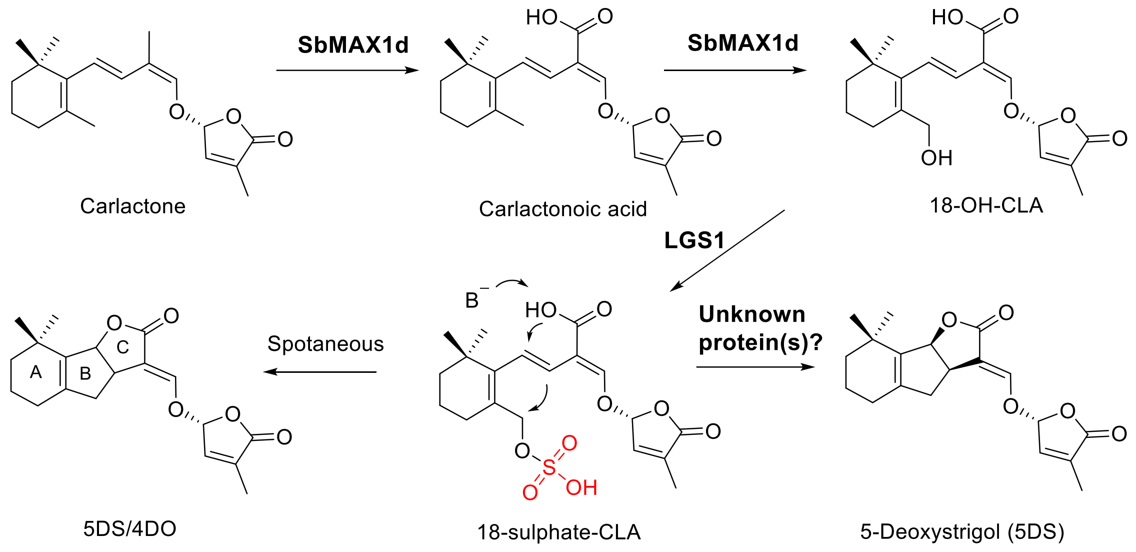


**Figure S8.** Proposed enzymic mechanism for the synthesis of 5DS and 4DO by SbMAX1d and LGS1. SbMAX1d can directly convert CL into 18-hydroxy-CLA. Then sulfotransferase LGS1 likely catalyze the sulfonation of C18 hydroxy to form the unstable intermediate 18-sulphate-CLA. The sulfonate group functions as a relatively easy leaving group, and under non-enzymatic conditions, the carboxylate trigger the $S_N2'$ type intramolecular nucleophilic attack and close the BC ring in a non-stereoselective manner, thereby generating a mixture of 4DO and 5DS. Although the nucleophilic attack step can be carried out spontaneously, there is likely a downstream enzyme catalyzes a more stereoselective conversion towards the synthesis of 5DS in Sorghum.

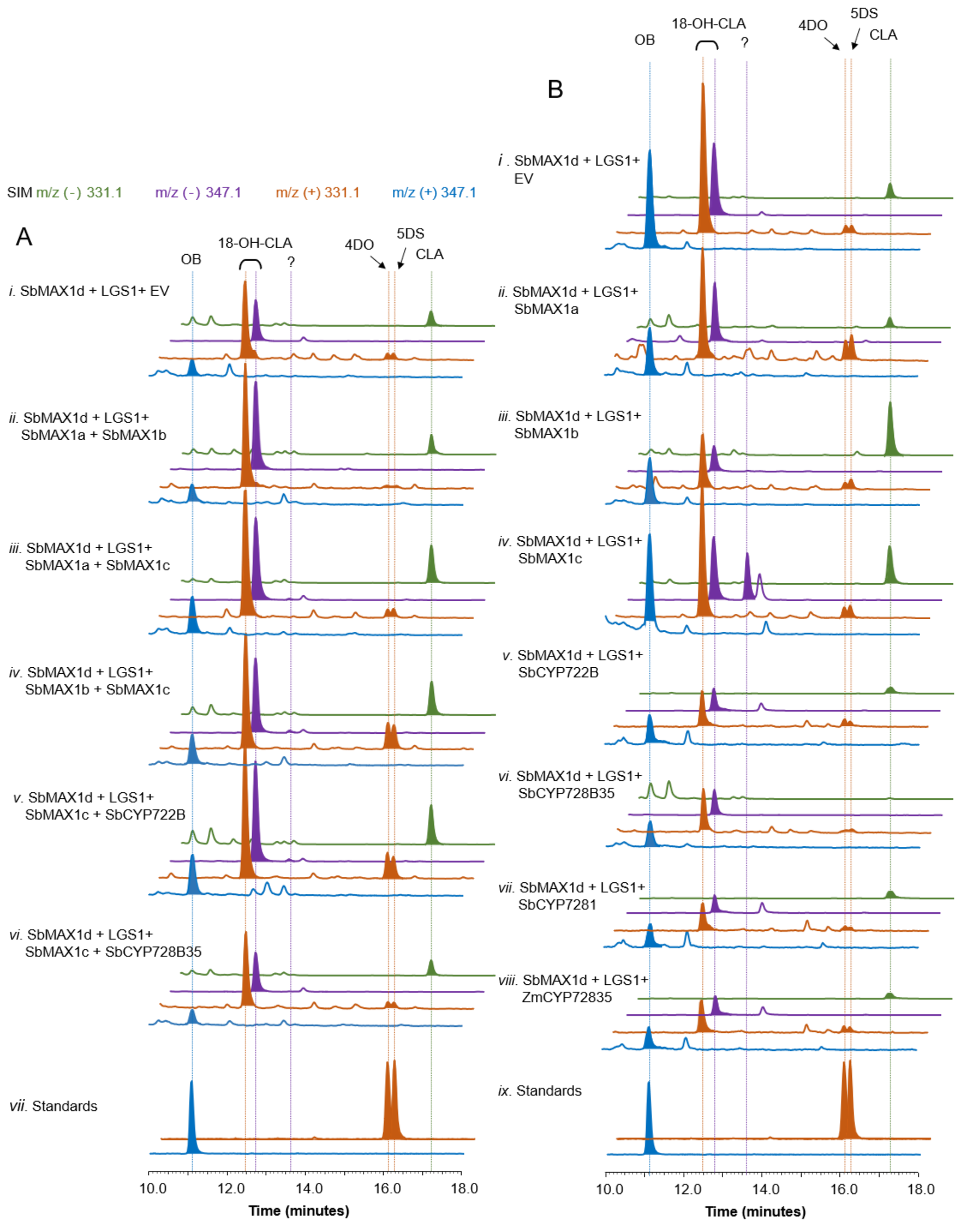


**Figure S9.** The enzymes after LGS1 and catalyze the exclusive conversion of 5DS is missing. (A) SIM EIC at m/z$^-$=331.1 (green), 347.1 (purple), and m/z$^+$=331.1 (orange), 347.1 (blue) of **ECL** (Table S2) cocultured with yeast strain SYL89 expressing SbMAX1d, LGS1 and i) EV (**YSL10N**, Table S2) ; ii) SbMAX1a and SbMAX1b; iii) SbMAX1a and SbMAX1c; iv) SbMAX1b and SbMAX1c; v) SbMAX1c and SbCYP722B; vi) SbMAX1c and SbCYP728B35 (**YSL10a-e**, Table S2); and vii) standards of OB, 4DO and 5DS. (B) SIM EIC at m/z$^-$=331.1 (green), 347.1 (purple), and m/z$^+$=331.1 (orange), 347.1 (blue) of **ECL** cocultured with yeast expressing ATR1, SbMAX1d, LGS1 and i) EV (**YSL9N**, Table S2), ii) SbMAX1a; iii) SbMAX1b; iv) SbMAX1c; v) SbCYP722B; vi) SbCYP728B35; vii) SbCYP728B1; viii) ZmCYP728B35 (**YSL9a-g**, Table S2) and ix) standards of OB, 4DO and 5DS.

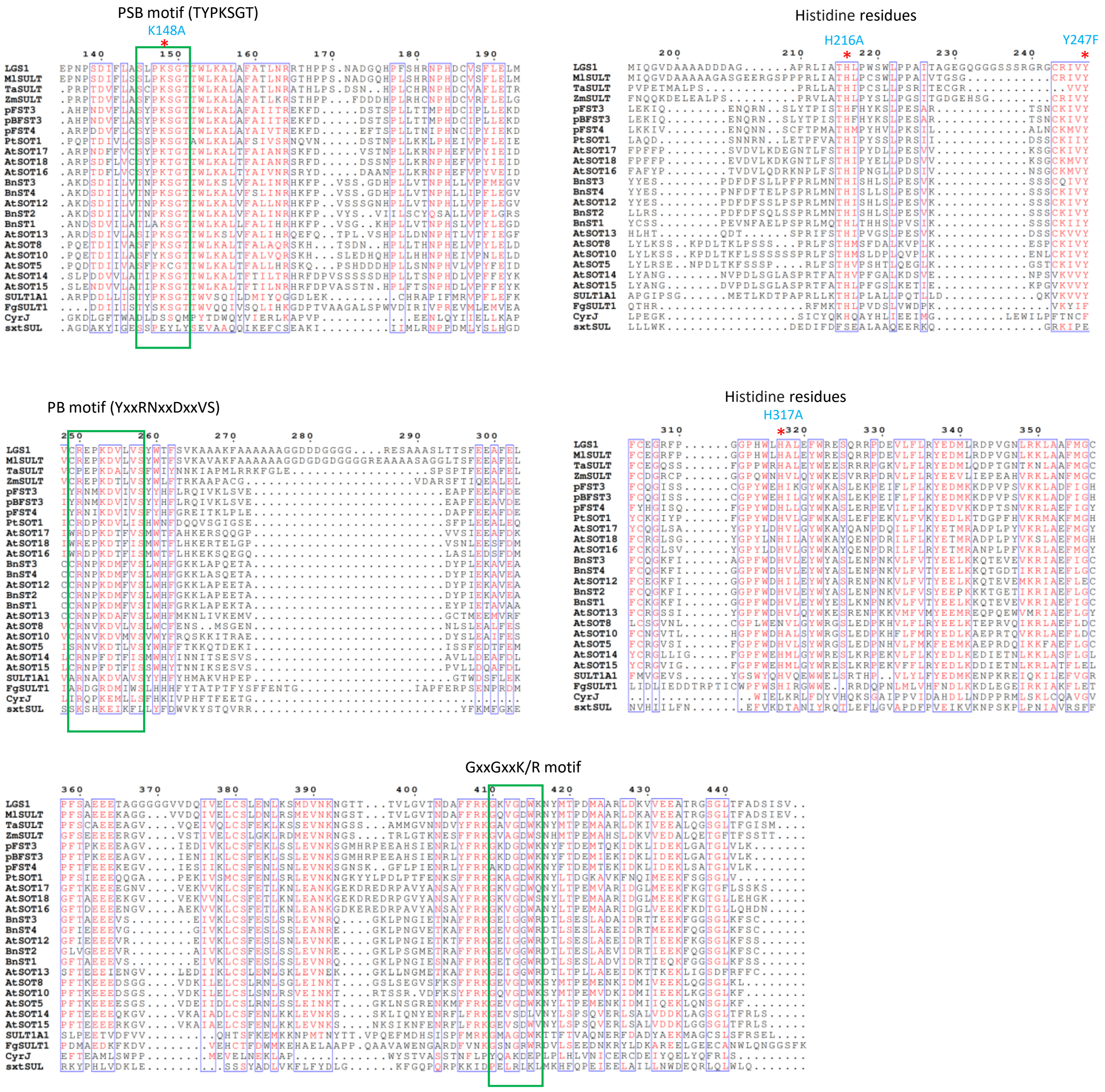


**Figure S10.** Full-length multiple sequence alignment of LGS1 with other SOTs. LGS1 was aligned with SOTs from plants, bacteria, and fungi using ClustalW (https://www.genome.jp/tools-bin/clustalw). The conserved PSB (TYPKSGT), PB (YxxRNxxDxxVS) and GxxGxxK/R motifs proposed to be critical to PAPS binding are highlighted with green box[2]. The mutated amino acid residues are marked with a red asterisk. The GenBank accession numbers of proteins analyzed are listed in Table S5.

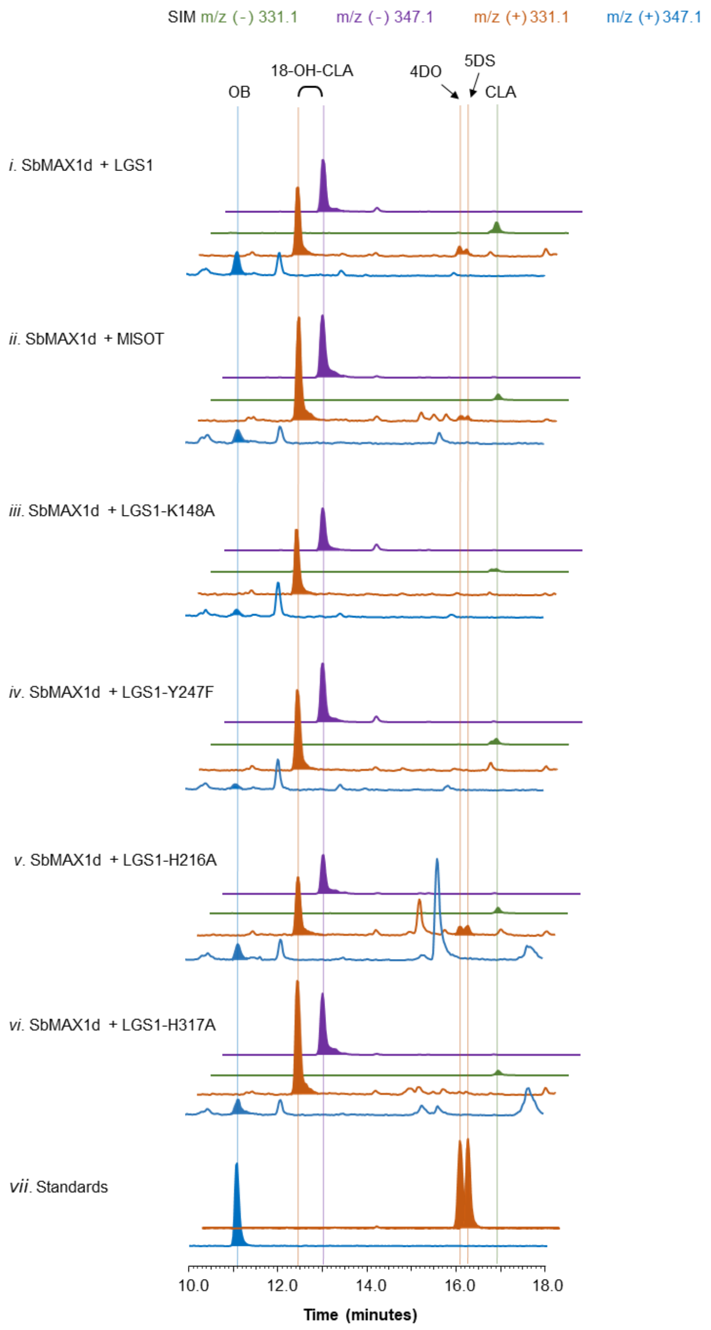


**Figure S11.** LC-MS analysis of LGS1 mutants and MlSOT. SIM EIC at $m/z^-$=331.1 (green), 347.1 (purple), and $m/z^+$=331.1 (orange), 347.1 (blue) of **ECL** (Table S2) cocultured with yeast expressing ATR1, SbMAX1d and i) LGS1 (**YSL8a**) ; ii) MlSOT (**YSL8e**); iii) LGS1$^{K148A}$ (**YSL8f**); iv) LGS1$^{Y247F}$ (**YSL8g**); v) LGS1$^{H216A}$ (**YSL8h**); vi) LGS1$^{H317A}$ (**YSL8i**); and vii) standards of OB, 4DO and 5DS. Strain information are detailed in Table S2.

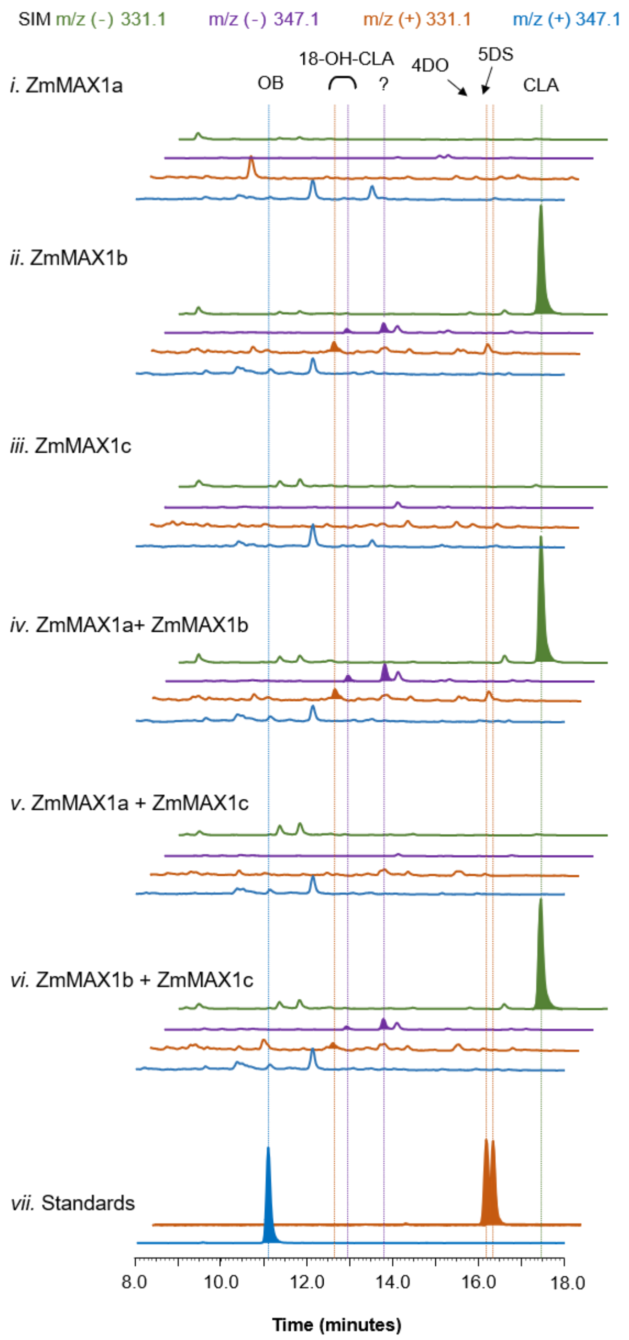


**Figure S12.** Functional characterization of MAX1 analogs from maize. SIM EIC at m/z$^-$=331.1 (green), 347.1 (purple), and m/z$^+$=331.1 (orange), 347.1 (blue) of **ECL** (Table S2) cocultured with yeast expressing ATR1, SbMAX1d, LGS1 and i) ZmMAX1a; ii) ZmMAX1b; iii) ZmMAX1c; iv) ZmMAX1a and ZmMAX1b; v) ZmMAX1a and ZmMAX1c; vi) ZmMAX1b and ZmMAX1c (**YSL11a-f**, Table S2); and vii) standards of OB, 4DO and 5DS.

**Table S1.** Plasmids used in the study.

| Plasmids | Description | Reference/ Source |
|---|---|---|
| pAC-BETAipi | Contains ctrE, crtB, crtI, crtY, and idi genes of *Erwinia herbicola* (Pantoea agglomerans) Eho10 and thereby produces beta-carotene in Escherichia coli; Derived from pACYCDuet-1, Replicon P15A (pACYC184); Resistance, Chloramphenicol; | Addgene #53277 [3] |
| pYL726 (pCDFDuet-trAtCCD7-OsD27) | pCDFDuet-1 carrying D27 from *Oryza sativa* and N-terminus 31 amino-acid truncated CCD7 from *Arabidopsis* | [4] |
| pYL735 (pET21a-trAtCCD8) | pET21a carrying N-terminus 56 amino-acid truncated CCD8 from *Arabidopsis* | [4] |
| pAG414GPD-ccdB | *Centromeric* TRP, attR1-$P_{GPD}$-*ccdB*-$T_{CYC1}$-attR2 | [5] |
| pAG415GPD-ccdB | *Centromeric* LEU, attR1-$P_{GPD}$-*ccdB*-$T_{CYC1}$-attR2 | [5] |
| pAG416GPD-ccdB | *Centromeric* URA, attR1-$P_{GPD}$-*ccdB*-$T_{CYC1}$-attR2 | [5] |
| pYL573 | *Centromeric* HIS, $P_{TEF1}$-*ATR1*-$T_{CYC1}$ | [6] |
| pYL759 | *Centromeric* LEU, $P_{PGK1}$-*AtMAX1*-$T_{pho5}$ | [4] |
| pYL777 | *Centromeric* TRP, $P_{GPD}$-*GaCYP722C* -$T_{CYC1}$ | [4] |
| pYL843 | *Centromeric* URA, $P_{GPD}$-*LGS1*-$T_{CYC1}$ | This study |
| pYL885 | *Centromeric* TRP, $P_{GPD}$-*SbMAX1b*-$T_{CYC1}$ | This study |
| pYL891 | *Centromeric* LEU, $P_{GPD}$-*SbMAX1d*-$T_{CYC1}$ | This study |
| pYL892 | *Centromeric* URA, $P_{GPD}$-*SbMAX1a*-$T_{CYC1}$ | This study |
| pYL893 | *Centromeric* URA, $P_{GPD}$-*SbMAX1c*-$T_{CYC1}$ | This study |
| pYL896 | *Centromeric* URA, $P_{GPD}$-*ZmSOT*-$T_{CYC1}$ | This study |
| pYL1051 | *Centromeric* URA, $P_{GPD}$-*TaSOT*-$T_{CYC1}$ | This study |
| pYL1052 | *Centromeric* URA, $P_{GPD}$-*ZmMAX1b*-$T_{CYC1}$ | This study |
| pYL1053 | *Centromeric* URA, $P_{GPD}$-*LGS1-2*-$T_{CYC1}$ | This study |
| pYL1061 | *Centromeric* URA, $P_{GPD}$-*SbMAX1b*-$T_{CYC1}$ | This study |
| pYL1063 | *Centromeric* TRP, $P_{GPD}$-*SbCYP722B*-$T_{CYC1}$ | This study |
| pYL1079 | *Centromeric* LEU, $P_{GPD}$- *ZmMAX1a*-$T_{CYC1}$ | This study |
| pYL1080 | *Centromeric* URA, $P_{GPD}$-*SbCYP728B35*-$T_{CYC1}$ | This study |
| pYL1084 | *Centromeric* LEU, $P_{GPD}$-*SbMAX1a*-$T_{CYC1}$ | This study |
| pYL1085 | *Centromeric* TRP, $P_{GPD}$-*ZmMAX1c*-$T_{CYC1}$ | This study |
| pYL1086 | *Centromeric* LEU, $P_{GPD}$-*ZmMAX1c*-$T_{CYC1}$ | This study |
| pYL1087 | *Centromeric* LEU, $P_{GPD}$-*ZmMAX1b*-$T_{CYC1}$ | This study |
| pYL1088 | *Centromeric* URA, $P_{GPD}$-*SbMAX1d*-$T_{CYC1}$ | This study |
| pYL1090 | *Centromeric* TRP, $P_{GPD}$-*LGS1*-$T_{CYC1}$ | This study |
| pYL1092 | *Centromeric* TRP, $P_{GPD}$-*SbMAX1c*-$T_{CYC1}$ | This study |
| pYL1095 | *Centromeric* HIS, $P_{GPD}$-*LGS1*-$T_{CYC1}$ | This study |
| pYL1097 | *Centromeric* URA, $P_{GPD}$-Zm*CYP728B35*-$T_{CYC1}$ | This study |
| pYL1102 | *Centromeric* URA, $P_{GPD}$-*SbCYP728B1*-$T_{CYC1}$ | This study |
| pYL1185 | *Centromeric* URA, $P_{GPD}$-*LGS1H216A mutant*-$T_{CYC1}$ | This study |
| pYL1192 | *Centromeric* URA, $P_{GPD}$-*MlSOT*-$T_{CYC1}$ | This study |
| pYL1243 | *Centromeric* URA, $P_{GPD}$-*LGS1Y247F mutant*-$T_{CYC1}$ | This study |
| pYL1246 | *Centromeric* URA, $P_{GPD}$-*LGS1K148A mutant*-$T_{CYC1}$ | This study |
| pYL1247 | *Centromeric* URA, $P_{GPD}$-*LGS1H317A mutant*-$T_{CYC1}$ | This study |

*P, promoter; T, terminator*

**Table S2.** Strains used in the study.

| Yeast Strain | Description | Genotype | Function | Reference |
|---|---|---|---|---|
| CEN.PK2-1D | | *MATα; his3D1; leu2-3_112; ura3-52; trp1-289; MAL2-8c; SUC2* | Wild type yeast strain | [7] |
| SYL89 | | CEN.PK2-1D, Leu2Δ::PTEF1-ATR1-TCYC1 | Yeast strain expressing the *A. thaliana* P450 reductase 1 | This study |
| YSL1 | CEN.PK2-1D carrying pYL573 and pYL759 | Centromeric HIS, $P_{TEF1}$-*ATR1*-$T_{CYC1}$, Centromeric URA, $P_{GPD}$-*AtMAX1*-$T_{CYC1}$ | CLA production | This study |
| YSL2a | CEN.PK2-1D carrying pYL573 and pYL892 | Centromeric HIS, $P_{TEF1}$-*ATR1*-$T_{CYC1}$, Centromeric URA, $P_{GPD}$-*SbMAX1a*-$T_{CYC1}$ | CLA production | This study |
| YSL2b | CEN.PK2-1D carrying pYL573, pYL885 and pAG416GPD-ccdB | Centromeric HIS, $P_{TEF1}$-*ATR1*-$T_{CYC1}$, Centromeric TRP, $P_{GPD}$-*SbMAX1b*-$T_{CYC1}$, Centromeric URA, $P_{GPD}$-*ccdB*-$T_{CYC1}$ | CLA production | This study |
| YSL2c | CEN.PK2-1D carrying pYL573 and pYL893 | Centromeric HIS, $P_{TEF1}$-*ATR1*-$T_{CYC1}$, Centromeric URA, $P_{GPD}$-*SbMAX1c*-$T_{CYC1}$ | CLA production | This study |
| YSL2d | CEN.PK2-1D carrying pYL573 and pYL891 | Centromeric HIS, $P_{TEF1}$-*ATR1*-$T_{CYC1}$, Centromeric LEU, $P_{GPD}$-*SbMAX1d*-$T_{CYC1}$ | OB and 18-OH-CLA production | This study |
| YSL3 | CEN.PK2-1D carrying pYL573 and pYL770 | Centromeric HIS, $P_{TEF1}$-*ATR1*-$T_{CYC1}$, Centromeric URA, $P_{PGK1}$-*OsCYP711A2*-$T_{PHO5}$ | 4DO production | [4] |
| YSL4 | CEN.PK2-1D carrying pYL573, pYL759, and pYL1070 | Centromeric HIS, $P_{TEF1}$-*ATR1*-$T_{CYC1}$, Centromeric LEU, $P_{PGK1}$-*AtMAX1*-$T_{PHO5}$, Centromeric TRP, $P_{GPD}$-*VuCYP722C*-$T_{CYC1}$ | OB production | [4] |
| YSL5 | CEN.PK2-1D carrying pYL573, pYL759, and pYL777 | Centromeric HIS, $P_{TEF1}$-*ATR1*-$T_{CYC1}$, Centromeric LEU, $P_{PGK1}$-*AtMAX1*-$T_{PHO5}$, Centromeric TRP, $P_{GPD}$-*GaCYP722C*-$T_{CYC1}$ | 5DS production | [4] |
| YSL6a | CEN.PK2-1D carrying pYL573, pYL891 and pYL885 | Centromeric HIS, $P_{TEF1}$-*ATR1*-$T_{CYC1}$, Centromeric LEU, $P_{GPD}$-*SbMAX1d*-$T_{CYC1}$, Centromeric TRP, $P_{GPD}$-*SbMAX1b*-$T_{CYC1}$, | OB and 18-OH-CLA production | This study |
| YSL6b | CEN.PK2-1D carrying pYL573, pYL891 and pYL892 | Centromeric HIS, $P_{TEF1}$-*ATR1*-$T_{CYC1}$, Centromeric LEU, $P_{GPD}$-*SbMAX1d*-$T_{CYC1}$, Centromeric URA, $P_{GPD}$-*SbMAX1a*-$T_{CYC1}$ | OB and 18-OH-CLA production | This study |
| YSL6c | CEN.PK2-1D carrying pYL573, pYL891 and pYL893 | Centromeric HIS, $P_{TEF1}$-*ATR1*-$T_{CYC1}$, Centromeric LEU, $P_{GPD}$-*SbMAX1d*-$T_{CYC1}$, Centromeric URA, $P_{GPD}$-*SbMAX1c*-$T_{CYC1}$ | OB ,18-OH-CLA and unknown production | This study |
| YSL6d | CEN.PK2-1D carrying pYL573, pYL885 and pYL892 | Centromeric HIS, $P_{TEF1}$-*ATR1*-$T_{CYC1}$, Centromeric TRP, $P_{GPD}$-*SbMAX1b*-$T_{CYC1}$, Centromeric URA, $P_{GPD}$-*SbMAX1a*-$T_{CYC1}$ | CLA production | This study |

| | | | | |
|---|---|---|---|---|
| YSL6e | CEN.PK2-1D carrying pYL573, pYL885 and pYL893 | Centromeric HIS, $P_{TEF1}$-*ATR1*-$T_{CYC1}$, Centromeric TRP, $P_{GPD}$-*SbMAX1b*-$T_{CYC1}$, Centromeric URA, $P_{GPD}$-*SbMAX1c*-$T_{CYC1}$ | CLA and unknown production | This study |
| YSL6f | CEN.PK2-1D carrying pYL573, pYL1084 and pYL893 | Centromeric HIS, $P_{TEF1}$-*ATR1*-$T_{CYC1}$, Centromeric LEU, $P_{GPD}$-*SbMAX1a*-$T_{CYC1}$, Centromeric URA, $P_{GPD}$-*SbMAX1c*-$T_{CYC1}$ | Unknown product | This study |
| YSL7N | CEN.PK2-1D carrying pYL573, pYL759, pYL777 and pAG416GPD-ccdB | Centromeric HIS, $P_{TEF1}$-*ATR1*-$T_{CYC1}$, Centromeric LEU, $P_{PGK1}$-*AtMAX1*-$T_{PHO5}$, Centromeric TRP, $P_{GPD}$-*GaCYP722C*-$T_{CYC1}$ Centromeric URA, $P_{GPD}$-*ccdB*-$T_{CYC1}$ | Negative control for YSL-6 | This study |
| YSL7a | CEN.PK2-1D carrying pYL573, pYL759, pYL777 and pYL892 | Centromeric HIS, $P_{TEF1}$-*ATR1*-$T_{CYC1}$, Centromeric LEU, $P_{PGK1}$-*AtMAX1*-$T_{PHO5}$, Centromeric TRP, $P_{GPD}$-*GaCYP722C*-$T_{CYC1}$ Centromeric URA, $P_{GPD}$-*SbMAX1a*-$T_{CYC1}$ | 5DS production, no downstream products detected | This study |
| YSL7b | CEN.PK2-1D carrying pYL573, pYL759, pYL777 and pYL1061 | Centromeric HIS, $P_{TEF1}$-*ATR1*-$T_{CYC1}$, Centromeric LEU, $P_{PGK1}$-*AtMAX1*-$T_{PHO5}$, Centromeric TRP, $P_{GPD}$-*GaCYP722C*-$T_{CYC1}$ Centromeric URA, $P_{GPD}$-*SbMAX1b*-$T_{CYC1}$ | 5DS production, no downstream products detected | This study |
| YSL7c | CEN.PK2-1D carrying pYL573, pYL759, pYL777 and pYL893 | Centromeric HIS, $P_{TEF1}$-*ATR1*-$T_{CYC1}$, Centromeric LEU, $P_{PGK1}$-*AtMAX1*-$T_{PHO5}$, Centromeric TRP, $P_{GPD}$-*GaCYP722C*-$T_{CYC1}$ Centromeric URA, $P_{GPD}$-*SbMAX1c*-$T_{CYC1}$ | 5DS production, no downstream products detected | This study |
| YSL7d | EN.PK2-1D carrying pYL573, pYL759, pYL777 and pYL1088 | Centromeric HIS, $P_{TEF1}$-*ATR1*-$T_{CYC1}$, Centromeric LEU, $P_{PGK1}$-*AtMAX1*-$T_{PHO5}$, Centromeric TRP, $P_{GPD}$-*GaCYP722C*-$T_{CYC1}$ Centromeric URA, $P_{GPD}$-*SbMAX1d*-$T_{CYC1}$ | 5DS production, no downstream products detected | This study |
| YSL8N | CEN.PK2-1D carrying pYL573, pYL891 and pAG416GPD-ccdB | Centromeric HIS, $P_{TEF1}$-*ATR1*-$T_{CYC1}$, Centromeric LEU, $P_{GPD}$-*SbMAX1d*-$T_{CYC1}$, Centromeric URA $P_{GPD}$-*ccdB*-$T_{CYC1}$ | Negative control for YSL-7 | This study |
| YSL8a | CEN.PK2-1D carrying pYL573, pYL891 and pYL843 | Centromeric HIS, $P_{TEF1}$-*ATR1*-$T_{CYC1}$, Centromeric LEU, $P_{GPD}$-*SbMAX1d*-$T_{CYC1}$, *Centromeric* URA, $P_{GPD}$-*LGS1*-$T_{CYC1}$ | LGS1 is involved in the conversion of 18-OH-CLA to 4DO/5DS | This study |
| YSL8b | CEN.PK2-1D carrying pYL573, pYL891 and pYL1053 | Centromeric HIS, $P_{TEF1}$-*ATR1*-$T_{CYC1}$, Centromeric LEU, $P_{GPD}$-*SbMAX1d*-$T_{CYC1}$, *Centromeric* URA, $P_{GPD}$-*LGS1v2*-$T_{CYC1}$ | LGS1-2 is involved in the conversion of 18-OH-CLA to 4DO/5DS | This study |
| YSL8c | CEN.PK2-1D carrying pYL573, pYL891 and pYL1051 | Centromeric HIS, $P_{TEF1}$-*ATR1*-$T_{CYC1}$, Centromeric LEU, $P_{GPD}$-*SbMAX1d*-$T_{CYC1}$, *Centromeric* URA, $P_{GPD}$-*TaSOT*-$T_{CYC1}$ | Failed functional characterization of TaSOT | This study |

| | | | | |
|---|---|---|---|---|
| YSL8d | CEN.PK2-1D carrying pYL573, pYL891 and pYL896 | Centromeric HIS, *P<sub></sub>*$P_{TEF1}$-*ATR1*-$T_{CYC1}$, Centromeric LEU, $P_{GPD}$-*SbMAX1d*-$T_{CYC1}$, *Centromeric* URA, $P_{GPD}$-*ZmSOT*-$T_{CYC1}$ | Failed functional characterization of ZmSOT | This study |
| YSL8e | CEN.PK2-1D carrying pYL573, pYL891 and pYL1192 | Centromeric HIS, $P_{TEF1}$-*ATR1*-$T_{CYC1}$, Centromeric LEU, $P_{GPD}$-*SbMAX1d*-$T_{CYC1}$, *Centromeric* URA, $P_{GPD}$-*MlSOT*-$T_{CYC1}$ | Trace 4DO/5DS production | This study |
| YSL8f | CEN.PK2-1D carrying pYL573, pYL891 and pYL1246 | Centromeric HIS, $P_{TEF1}$-*ATR1*-$T_{CYC1}$, Centromeric LEU, $P_{GPD}$-*SbMAX1d*-$T_{CYC1}$, *Centromeric* URA, $P_{GPD}$- *LGS1K148A mutant* -$T_{CYC1}$ | No 4DO/5DS production | This study |
| YSL8g | CEN.PK2-1D carrying pYL573, pYL891 and pYL1243 | Centromeric HIS, $P_{TEF1}$-*ATR1*-$T_{CYC1}$, Centromeric LEU, $P_{GPD}$-*SbMAX1d*-$T_{CYC1}$, *Centromeric* URA, $P_{GPD}$- *LGS1Y247F mutant* -$T_{CYC1}$ | No 4DO/5DS production | This study |
| YSL8h | CEN.PK2-1D carrying pYL573, pYL891 and pYL1185 | Centromeric HIS, $P_{TEF1}$-*ATR1*-$T_{CYC1}$, Centromeric LEU, $P_{GPD}$-*SbMAX1d*-$T_{CYC1}$, *Centromeric* URA, $P_{GPD}$- *LGS1H216A mutant* -$T_{CYC1}$ | 4DO/5DS production | This study |
| YSL8i | CEN.PK2-1D carrying pYL573, pYL891 and pYL1247 | Centromeric HIS, $P_{TEF1}$-*ATR1*-$T_{CYC1}$, Centromeric LEU, $P_{GPD}$-*SbMAX1d*-$T_{CYC1}$, *Centromeric* URA, $P_{GPD}$- *LGS1H317A mutant* -$T_{CYC1}$ | No 4DO/5DS production | This study |
| YSL9N | CEN.PK2-1D carrying pYL573, pYL891, pYL1090 and pAG416GPD-ccdB | Centromeric HIS, $P_{TEF1}$-*ATR1*-$T_{CYC1}$, Centromeric LEU, $P_{GPD}$-*SbMAX1d*-$T_{CYC1}$, Centromeric TRP, $P_{GPD}$-*LGS1*-$T_{CYC1}$, Centromeric URA, $P_{GPD}$-*ccdB*-$T_{CYC1}$ | Negative control for YSL-8 | This study |
| YSL9a | CEN.PK2-1D carrying pYL573, pYL891 pYL1090 and pYL892 | Centromeric HIS, $P_{TEF1}$-*ATR1*-$T_{CYC1}$, Centromeric LEU, $P_{GPD}$-*SbMAX1d*-$T_{CYC1}$, Centromeric TRP, $P_{GPD}$-*LGS1*-$T_{CYC1}$, Centromeric URA, $P_{GPD}$-*SbMAX1a*-$T_{CYC1}$ | No change in 4DO/5DS ratio or new products detected | This study |
| YSL9b | CEN.PK2-1D carrying pYL573, pYL891, pYL885 and pYL843 | Centromeric HIS, $P_{TEF1}$-*ATR1*-$T_{CYC1}$, Centromeric LEU, $P_{GPD}$-*SbMAX1d*-$T_{CYC1}$, Centromeric TRP, $P_{GPD}$-*SbMAX1b*-$T_{CYC1}$, Centromeric URA, $P_{GPD}$-*LGS1*-$T_{CYC1}$ | No change in 4DO/5DS ratio or new products detected | This study |
| YSL9c | CEN.PK2-1D carrying pYL573, pYL891, pYL1090 and pYL893 | Centromeric HIS, $P_{TEF1}$-*ATR1*-$T_{CYC1}$, Centromeric LEU, $P_{GPD}$-*SbMAX1d*-$T_{CYC1}$, Centromeric TRP, $P_{GPD}$-*LGS1*-$T_{CYC1}$, Centromeric URA, $P_{GPD}$-*SbMAX1c*-$T_{CYC1}$ | No change in 4DO/5DS ratio or new products detected | This study |

| | | | | |
|---|---|---|---|---|
| YSL9d | CEN.PK2-1D carrying pYL573, pYL891, pYL843 and pYL1063 | Centromeric HIS, $P_{TEF1}$-*ATR1*-$T_{CYC1}$, Centromeric LEU, $P_{GPD}$-*SbMAX1d*-$T_{CYC1}$, Centromeric URA, $P_{GPD}$-*LGS1*-$T_{CYC1}$, Centromeric TRP, $P_{GPD}$-*SbCYP722B*-$T_{CYC1}$ | No change in 4DO/5DS ratio or new products detected | This study |
| YSL9e | CEN.PK2-1D carrying pYL573, pYL891, pYL1090 and pYL1080 | Centromeric HIS, $P_{TEF1}$-*ATR1*-$T_{CYC1}$, Centromeric LEU, $P_{GPD}$-*SbMAX1d*-$T_{CYC1}$, Centromeric TRP, $P_{GPD}$-*LGS1*-$T_{CYC1}$, Centromeric URA, $P_{GPD}$-*SbCYP728B35*-$T_{CYC1}$ | No change in 4DO/5DS ratio or new products detected | This study |
| YSL9f | CEN.PK2-1D carrying pYL573, pYL891, pYL1090 and pYL1102 | Centromeric HIS, $P_{TEF1}$-*ATR1*-$T_{CYC1}$, Centromeric LEU, $P_{GPD}$-*SbMAX1d*-$T_{CYC1}$, Centromeric TRP, $P_{GPD}$-*LGS1*-$T_{CYC1}$, Centromeric URA, $P_{GPD}$-*SbCYP728B1*-$T_{CYC1}$ | No change in 4DO/5DS ratio or new products detected | This study |
| YSL9g | CEN.PK2-1D carrying pYL573, pYL891, pYL1090 and pYL1097 | Centromeric HIS, $P_{TEF1}$-*ATR1*-$T_{CYC1}$, Centromeric LEU, $P_{GPD}$-*SbMAX1d*-$T_{CYC1}$, Centromeric TRP, $P_{GPD}$-*LGS1*-$T_{CYC1}$, Centromeric URA, $P_{GPD}$-*ZmCYP728B35*-$T_{CYC1}$ | No change in 4DO/5DS ratio or new products detected | This study |
| YSL10N | SYL89 carrying pYL1095, pYL891, pAG414GPD-ccdB and pAG416GPD-ccdB | Centromeric HIS, $P_{GPD}$-*LGS1*-$T_{CYC1}$, Centromeric LEU, $P_{GPD}$-*SbMAX1d*-$T_{CYC1}$, Centromeric TRP, $P_{GPD}$-*ccdB*-$T_{CYC1}$, Centromeric URA, $P_{GPD}$-*ccdB*-$T_{CYC1}$ | No change in 4DO/5DS ratio or new products detected | This study |
| YSL10a | SYL89 carrying pYL1095, pYL891, pYL885 and pYL892 | Centromeric HIS, $P_{GPD}$-*LGS1*-$T_{CYC1}$, Centromeric LEU, $P_{GPD}$-*SbMAX1d*-$T_{CYC1}$, Centromeric TRP, $P_{GPD}$-*SbMAX1b*-$T_{CYC1}$, Centromeric URA, $P_{GPD}$-*SbMAX1a*-$T_{CYC1}$ | No change in 4DO/5DS ratio or new products detected | This study |
| YSL10b | SYL89 carrying pYL1095, pYL891, pYL1092 and pYL892 | Centromeric HIS, $P_{GPD}$-*LGS1*-$T_{CYC1}$, Centromeric LEU, $P_{GPD}$-*SbMAX1d*-$T_{CYC1}$, Centromeric TRP, $P_{GPD}$-*SbMAX1c*-$T_{CYC1}$, Centromeric URA, $P_{GPD}$-*SbMAX1a*-$T_{CYC1}$ | No change in 4DO/5DS ratio or new products detected | This study |
| YSL10c | SYL89 carrying pYL1095, pYL891, pYL885 and pYL893 | Centromeric HIS, $P_{GPD}$-*LGS1*-$T_{CYC1}$, Centromeric LEU, $P_{GPD}$-*SbMAX1d*-$T_{CYC1}$, Centromeric TRP, $P_{GPD}$-*SbMAX1b*-$T_{CYC1}$, Centromeric URA, $P_{GPD}$-*SbMAX1c* -$T_{CYC1}$ | No change in 4DO/5DS ratio or new products detected | This study |
| YSL10d | SYL89 carrying pYL1095, pYL891, pYL1063 and pYL893 | Centromeric HIS, $P_{GPD}$-*LGS1*-$T_{CYC1}$, Centromeric LEU, $P_{GPD}$-*SbMAX1d*-$T_{CYC1}$, Centromeric TRP, $P_{GPD}$-*SbCYP722B*-$T_{CYC1}$, Centromeric URA, $P_{GPD}$-*SbMAX1c*-$T_{CYC1}$ | No change in 4DO/5DS ratio or new products detected | This study |

| | | | | |
|---|---|---|---|---|
| YSL10e | SYL89 carrying pYL1095, pYL891, pYL1092 and pYL1080 | Centromeric HIS, $P_{GPD}$-*LGS1*-$T_{CYC1}$, Centromeric LEU, $P_{GPD}$-*SbMAX1d*-$T_{CYC1}$, Centromeric TRP, $P_{GPD}$-*SbMAX1c*-$T_{CYC1}$, Centromeric URA, $P_{GPD}$-*SbCYP728B35*-$T_{CYC1}$ | No change in 4DO/5DS ratio or new products detected | This study |
| YSL11a | CEN.PK2-1D carrying pYL573 and pYL1079 | Centromeric HIS, $P_{TEF1}$-*ATR1*-$T_{CYC1}$, *Centromeric* LEU, $P_{GPD}$-*ZmMAX1a*-$T_{CYC1}$ | Failed functional characterization of ZmMAX1a | This study |
| YSL11b | CEN.PK2-1D carrying pYL573 and pYL1087 | Centromeric HIS, $P_{TEF1}$-*ATR1*-$T_{CYC1}$, *Centromeric* LEU, $P_{GPD}$-*ZmMAX1b*-$T_{CYC1}$ | CLA production | This study |
| YSL11c | CEN.PK2-1D carrying pYL573 and pYL1086 | Centromeric HIS, $P_{TEF1}$-*ATR1*-$T_{CYC1}$, *Centromeric* LEU, $P_{GPD}$-*ZmMAX1c*-$T_{CYC1}$ | Failed functional characterization of ZmMAX1c | This study |
| YSL11d | CEN.PK2-1D carrying pYL573, pYL1079 and pYL1052 | Centromeric HIS, $P_{TEF1}$-*ATR1*-$T_{CYC1}$, *Centromeric* LEU, $P_{GPD}$-*ZmMAX1a*-$T_{CYC1}$, *Centromeric* URA, $P_{GPD}$-*ZmMAX1b*-$T_{CYC1}$ | CLA production | This study |
| YSL11e | CEN.PK2-1D carrying pYL573, pYL1079 and pYL1085 | Centromeric HIS, $P_{TEF1}$-*ATR1*-$T_{CYC1}$, *Centromeric* LEU, $P_{GPD}$-*ZmMAX1a*-$T_{CYC1}$, *Centromeric* TRP, $P_{GPD}$-*ZmMAX1c*-$T_{CYC1}$ | Failed functional characterization of ZmMAX1a/ ZmMAX1c | This study |
| YSL11f | CEN.PK2-1D carrying pYL573, pYL1087 and pYL1085 | Centromeric HIS, $P_{TEF1}$-*ATR1*-$T_{CYC1}$, *Centromeric* LEU, $P_{GPD}$-*ZmMAX1b*-$T_{CYC1}$, *Centromeric* TRP, $P_{GPD}$-*ZmMAX1c*-$T_{CYC1}$ | CLA production | This study |
| ***E. coli* Strain** | **Base Strain** | **Plasmid** | **Function** | |
| ECL | BL21(DE3) | pAC-BETAipi; pYL726 (pCDFDuet-trAtCCD7-OsD27); pYL735 (pET21a-trAtCCD8) | CL production | [4] |

**Table S3.** CYPs used in this study and summary of results. The accession numbers of amino acid sequences of sorghum genes were extracted from Phytozome database (*Sorghum bicolor v3.1.1*; https://phytozome-next.jgi.doe.gov/), the others are from NCBI.

| Gene | Species | Notation | Experimental results/product | Ref |
|---|---|---|---|---|
| *AtMAX1* (AK316903) | *Arabidopsis thaliana* | CL → CLA | CLA | 8, 9 |
| *SbMAX1a* (Sobic.010G170400) | *Sorghum bicolor* | N.D. | CL → CLA, weak | 10 |
| *SbMAX1b* (Sobic.003G269600) | *Sorghum bicolor* | CL → CLA | CL → CLA | 10 |
| *SbMAX1c* (Sobic.004G095500) | *Sorghum bicolor* | N.D. | CL → CLA & U.P. | 10 |
| *SbMAX1d* (Sobic.003G269500) | *Sorghum bicolor* | N.D. | CL → 18-OH-CLA→OB | 10 |
| *ZmMAX1a* (PWZ07057) | *Sorghum bicolor* | CL → CLA, weak | N.D. | 8 |
| *ZmMAX1b* (ONM29770) | *Sorghum bicolor* | CL → CLA & 4DO → OB | CL → CLA, trace 18-OH-CLA & U.P. | 8 |
| *ZmMAX1c* (XP_020407074) | *Sorghum bicolor* | CL → CLA, weak | N.D. | 8 |
| *SbCYP722B* (Sobic.009G000700) | *Sorghum bicolor* | | N.D. | |
| *SbCYP728B35* (Sobic.008G122800) | *Sorghum bicolor* | | N.D. | 10 |
| *SbCYP728B1* (Sobic.002G336100) | *Sorghum bicolor* | | N.D. | 10 |
| *ZmCYP728B35* (NP_001148166) | *Zea mays* | | N.D. | |

U.P. Unknown product; N.D. No detected; ZmMAX1a variation used in this study (PWZ07057) is nine amino acids (ASDTTMQRH) longer than ZmMAX1a (FJ957947) reported before[8]; ZmMAX1c variation (XP_020407074) used here is one amino acid (Q) longer than ZmMAX1c (NP_001145812) reported before[8].

**Table S4.** Accession numbers of MAX1 analogs used for the phylogenetic tree analysis in Figure 2 and Figure S1. The amino acid sequences are downloadable from NCBI, except for SfMAX1, which is downloaded from https://phytozome.jgi.doe.gov/pz/portal.html.

| Gene | Species | Size (a.a.) | Accession numbers |
|---|---|---|---|
| *AtMAX1* | *Arabidopsis thaliana* | 522 | NP_565617 |
| *PtMAX1a* | *Populus trichocarpa* | 529 | XP_006372016 |
| *PtMAX1b* | *Populus trichocarpa* | 529 | XP_006382011 |
| *PhMAX1* | *Petunia x hybrida* | 533 | AEB97383 |
| *SmMAX1a* | *Selaginella moellendorffii* | 512 | AGI65366 |
| *SmMAX1b* | *Selaginella moellendorffii* | 512 | BBA85738 |
| *OsCYP711A2* (*Os01g0700900*) | *Oryza sativa* (*rice*) | 539 | XP_015633367 |
| *OsCYP711A3* (*Os01g0701400*) | *Oryza sativa* (*rice*) | 541 | XP_015644699 |
| *OsCYP711A4* (*Os01g0701500*) | *Oryza sativa* (*rice*) | 516 | XP_015642272 |
| *OsCYP711A5* (*Os02g0221900*) | *Oryza sativa* (*rice*) | 548 | XP_015626073 |
| *OsCYP711A6* (*Os06g0565100*) | *Oryza sativa* (*rice*) | 540 | XP_015644019 |
| *SlMAX1* | *Solanum lycopersicum* (*tomato*) | 519 | XP_004245085 |
| *StMAX1* | *Solanum tuberosum* | 519 | XP_006351579 |
| *ZmMAX1a* | *Zea mays* | 537 | PWZ07057 |
| *ZmMAX1b* | *Zea mays* | 543 | ONM29770 |
| *ZmMAX1c* | *Zea mays* | 560 | XP_020407074 |
| *AmtMAX1* | *Amborella trichopoda* | 555 | XP_011626843 |
| *AcMAX1* | *Aquilegia coerulea* (*columbine*) | 539 | PIA42995 |
| *AlMAX1* | *Arabidopsis lyrata* | 522 | XP_020884785 |
| *BrMAX1* | *Brassica rapa* | 530 | RID70458 |
| *CcMAX1a* | *Cajanus cajanifolius* | 531 | XP_020207800 |
| *CcMAX1b* | *Cajanus cajanifolius* | 534 | XP_020210968 |
| *CsMAX1* | *Cannabis sativa* | 525 | XP_030490461 |
| *CrMAX1* | *Capsella rubella* | 525 | XP_006294008 |
| *CaaMAX1* | *Capsicum annuum* | 532 | XP_016537773 |
| *CaMAX1a* | *Cicer arietinum* | 527 | XP_004498873 |
| *CaMAX1b* | *Cicer arietinum* | 543 | XP_004501462 |
| *CisMAX1* | *Citrus sinensis* | 547 | XP_006466891 |
| *CcMAX1a* | *Citrus clementina* | 538 | XP_006450866 |
| *CcMAX1b* | *Citrus clementina* | 547 | XP_006425560 |
| *CusMAX1* | *Cucumis sativus* | 529 | XP_004141322 |
| *EgMAX1* | *Eucalyptus grandis* | 542 | XP_010047358 |
| *EsMAX1* | *Eutrema salsugineum* | 527 | XP_006408861 |
| *FvMAX1* | *Fragaria vesca* | 531 | XP_004291053 |
| *GrMAX1* | *Gossypium raimondii* | 539 | XP_012442761 |
| *LjMax1* | *Lotus japonicus* | 538 | BBM90835 |
| *NnMAX1* | *Nelumbo nucifera* (*sacred lotus*) | 544 | XP_010262061 |
| *MeMAX1* | *Manihot esculenta* | 527 | XP_021598065 |
| *EgMAX1* | *Erythranthe guttata* | 526 | XP_012857912 |
| *MaMAX1a* | *Musa acuminata* | 529 | XP_009380454 |
| *MaMAX1b* | *Musa acuminata* | 526 | XP_009408870 |
| *ObMAX1* | *Oryza brachyantha* | 543 | XP_006646245 |

| | | | |
|---|---|---|---|
| *PdMAX1a* | *Phoenix dactylifera* | 529 | XP_008793212 |
| *PdMAX1b* | *Phoenix dactylifera* | 545 | XP_008782508 |
| *PmMAX1a* | *Prunus mume* | 538 | XP_008220493 |
| *PmMAX1b* | *Prunus mume* | 541 | XP_008220494 |
| *PpMAX1a* | *Prunus persica* | 538 | XP_007222310 |
| *PpMAX1b* | *Prunus persica* | 533 | XP_007224581 |
| *PpMAX1c* | *Prunus persica* | 536 | XP_007225050 |
| *MdMAX1a* | *Malus domestica* | 526 | XP_008357300 |
| *MdMAX1b* | *Malus domestica* | 542 | RXH72971 |
| *MdMAX1c* | *Malus domestica* | 540 | XP_008393629 |
| *MdMAX1d* | *Malus domestica* | 532 | XP_028955031 |
| *VvMAX1* | *Vitis vinifera* | 530 | XP_002279086 |
| *CpMAX1* | *Carica papaya* | 538 | XP_021907713 |
| *RcMAX1* | *Ricinus communis* | 540 | XP_002516084 |
| *SiMAX1a* | *Setaria italica* | 533 | XP_004969659 |
| *SiMAX1b* | *Setaria italica* | 536 | XP_004969660 |
| *SiMAX1c* | *Setaria italica* | 549 | XP_004951458 |
| *GmMAX1a* | *Glycine max* | 551 | AQY54419 |
| *GmMAX1b* | *Glycine max* | 548 | AQY54420 |
| *GmMAX1c* | *Glycine max* | 532 | XP_003549345 |
| *GmMAX1d* | *Glycine max* | 538 | XP_003544542 |
| *PgMAX1* | *Picea glauca* | 544 | AGI65359 |
| *SaMAX1* | *Sinapis alba* | 529 | KAF8102838 |
| *TcMAX1* | *Theobroma cacao* | 539 | XP_007012311 |
| *CmMAX1* | *Cucurbita moschata* | 531 | XP_022936422 |
| *TtMAX1a* | *Triticum turgidum subsp. durum* | 536 | VAH72491 |
| *TtMAX1b* | *Triticum turgidum subsp. durum* | 530 | VAI09923 |
| *TaMAX1a* | *Triticum aestivum* | 518 | KAF7096775 |
| *TaMAX1b* | *Triticum aestivum* | 534 | KAF7025803 |
| *TaMAX1c* | *Triticum aestivum* | 518 | KAF7102706 |
| *TaMAX1d* | *Triticum aestivum* | 536 | KAF6994362 |
| *TaMAX1e* | *Triticum aestivum* | 530 | KAF7045862 |
| *TaMAX1f* | *Triticum aestivum* | 530 | KAF7102157 |
| *TaMAX1g* | *Triticum aestivum* | 516 | KAF7041259 |
| *AetMAX1* | Aegilops tauschii | 529 | XP_020163444 |
| *BdMAX1a* | *Brachypodium distachyon* | 530 | XP_003560652 |
| *BdMAX1b* | *Brachypodium distachyon* | 528 | XP_003575594 |
| *BdMAX1c* | *Brachypodium distachyon* | 531 | XP_003571126 |
| *BdMAX1d* | *Brachypodium distachyon* | 525 | XP_010237353 |
| *BdMAX1e* | *Brachypodium distachyon* | 534 | XP_003562092 |
| *HvMAX1a* | *Hordeum vulgare* | 524 | KAE8788859 |
| *HvMAX1b* | *Hordeum vulgare* | 533 | KAE8810993 |
| *HvMAX1c* | *Hordeum vulgare* | 537 | BAJ97619 |
| *HvMAX1d* | *Hordeum vulgare* | 562 | KAE8781561 |
| *HvMAX1e* | *Hordeum vulgare* | 526 | KAE8781562 |
| *MtMAX1a* | *Medicago truncatula* | 529 | AGI65361 |
| *MtMAX1b* | *Medicago truncatula* | 541 | AGI65360 |
| *SbMAX1a* | *Sorghum bicolor* | 540 | XP_002438586 |
| *SbMAX1b* | *Sorghum bicolor* | 545 | XP_002456213 |
| *SbMAX1c* | *Sorghum bicolor* | 545 | XP_002453551 |
| *SbMAX1d* | *Sorghum bicolor* | 547 | XP_002458367 |
| *MlMAX1a* | *Miscanthus lutarioriparius* | 528 | CAD6336534 |
| *MlMAX1b* | *Miscanthus lutarioriparius* | 521 | CAD6237234 |

| | | | |
|---|---|---|---|
| *MlMAX1c* | *Miscanthus lutarioriparius* | 540 | CAD6247547 |
| *MlMAX1d* | *Miscanthus lutarioriparius* | 539 | CAD6237236 |
| *KnMAX1a* | *Klebsormidium nitens* | 537 | GAQ86604 |
| *KnMAX1b* | *Klebsormidium nitens* | 569 | GAQ88536 |
| *KnMAX1c* | *Klebsormidium nitens* | 574 | GAQ88955 |
| *KnMAX1d* | *Klebsormidium nitens* | 550 | GAQ88956 |
| *SfMAX1* | *Sphagnum fallax* | 503 | Sphfalx0047s0130 |

**Table S5.** Accession numbers of SOTs used for the phylogenetic analysis in Figure 3. The amino acid sequences are downloadable from NCBI. The functions of some plant SOTs have been identified before[11].

| Gene | Species | Size (a.a.) | Accession numbers | Reference |
|---|---|---|---|---|
| *AtSOT13* | *Arabidopsis thaliana* | 324 | NP_178472 | 12 |
| *AtSOT8* | *Arabidopsis thaliana* | 331 | NP_172799 | 13 |
| *AtSOT14* | *Arabidopsis thaliana* | 347 | NP_196317 | 14 |
| *AtSOT15* | *Arabidopsis thaliana* | 359 | NP_568177 | 14 |
| *AtSOT5* | *Arabidopsis thaliana* | 323 | NP_190093 | 12 |
| *pFST3* | *Flaveria chlorifolia* | 312 | P52836 | 15 |
| *pFST4'* | *Flaveria chlorifolia* | 320 | P52837 | 15 |
| *AtSOT10* | *Arabidopsis thaliana* | 333 | NP_179098 | 16 |
| *AtSOT12* | *Arabidopsis thaliana* | 326 | NP_178471 | 16 |
| *AtSOT16* | *Arabidopsis thaliana* | 338 | NP_177550 | 17 |
| *AtSOT17* | *Arabidopsis thaliana* | 346 | NP_173294 | 17 |
| *AtSOT18* | *Arabidopsis thaliana* | 350 | NP_177549 | 17 |
| *AtTPST* | *Arabidopsis thaliana* | 500 | NP_001320287 | 18 |
| *PtSOT1* | *Populus trichocarpa* | 350 | XP_002318367 | 19 |
| *pBFST3* | *Flaveria bidentis* | 312 | P52835 | 20 |
| *BnST1* | *Brassica napus* | 323 | XP_013723132 | 21 |
| *BnST2* | *Brassica napus* | 324 | AAC63112 | 21 |
| *BnST3* | *Brassica napus* | 325 | XP_013606782 | 21 |
| *BnST4* | *Brassica napus* | 323 | NP_001302500 | 21, 22 |
| *LGS1* | *Sorghum bicolor* | 452 | KAG0530922 | 23 |
| *MlSOT* | *Miscanthus lutarioriparius* | 401 | CAD6255761 | |
| *TaSOT* | *Triticum aestivum* | 345 | KAF7005357 | |
| *ZmSOT* | *Zea mays* | 451 | XP_008672387 | |
| *SxtSULT* | *Microseira wollei* | 302 | ACG63834 | 24 |
| *CyrJ* | *Cylindrospermopsis raciborskii* | 261 | WP_007357796 | 25 |
| *FgSULT1* | *Fusarium graminearum* PH-1 | 307 | XP_011322917 | 2 |
| *FvSULT* | *Fusarium verticillioides* | 309 | RBR08306 | 2 |
| *XlSULT* | *Xylaria longipes* | 553 | RYC60899 | 2 |
| *SULT1A1* | *Homo sapiens* | 295 | NP_001046 | 2 |
| *HsSULT* | *Homo sapiens* | 411 | NP_004798 | 2 |
| *MmSULT* | *Mus musculus* | 411 | NP_056633 | 2 |

**Table S6.** Sequences of genes used in this study

| Gene | Sequence (5′–3′) |
|---|---|
| *SbMAX1a* | ATGGAAATGGCTGGTGCTGCTGGTACTGCTGAAACTTGGTTGCCATATGTTACTA<br>CTGCTGCTTCTTGTGCTGTTGCTGTTTTTTTCTTGTTGTACTTCTATGCTCCACAAT<br>GGGCTGTTAGAGGTGTTCCAGGTCCACCAGCTTTGCCAGTTGTTGGTCATTTGCC<br>ATTATTGGCTAGACATGGTCCAGATATTTTTGGTTTGTTGGCTAAAAGTACGGCC<br>CAATCTTTAGATTCCATTTGGGTAGACAACCATTGGTTATAGTTGCTGATCCAGAA<br>TTGTGTAGAGAAGTTGGTGTTAGACAGTTCAAGTTGATCCCAAATAGATCTTTGCC<br>AGCTCCAATTGCTGGTTCTCCATTGCATCAAAAGGGTTTGTTTTTTACCTCCAGAG<br>ATGAGAGATGGTCTGCTATGAGAAACACCATCATTAGCTTGTACCAACCATCTCAT<br>TTGGCTGGTTTGGTTCCAACTATGCAAAGATGTATTGAAAGAGCTGCTGACGCTA<br>TTTTGGCTCCAGGTGTTCAACAAAATGGTGATGGTGATGTTGACGTCGATGTTGA<br>TTTTTCCGATCTGTCTTTGAAGTTGGCCACCGATATTATTGGTCAAGCTGCTTTTG<br>GTGTTGATTTCGGTTTGACTGCTTCTGGTGATCCAGGTGGTGAAGCTGCTGAATT<br>CATTAGAGAACACGTTCATTCTACCACCTCATTGAAGATGGATTTGTCTGCTCCAT<br>TGTCTGTTGCTTTGGGTTTAGTTGCTCCAGCTTTACAAGGTCCAGTTAGAAGATTA<br>TTGTCTAGAGTTCCAGGTACAGCTGATTGGAAAGTTGCTAGAACTAATGCTAGATT<br>GAGAGCCAGAGTTGATGAAGTTGTTGCTGCTAGAGCAAGAGCTAGAGAAAGACG<br>TAGACACGGTGAAGCTAGAACAAAGGATTTTTTGTCAGCTGTTTTGGATGCCAGA<br>GATAGATCTGCTGCTTTGAGAGAATTATTGACCCCAGATCATGTTTCTGCTTTGAC<br>CTATGAACATTTGTTAGCTGGTTCTGCTACTACCGCTTTTACTTTATCTTCTGCCGT<br>TTATTTGGTTGCCGGTCATCCAGAAGTTGAAGCTAAGTTGTTGGCTGAAGTTGAT<br>GGTTTTGGTCCAAGAGGTGCTGTTCCAACTGCTGATGACTTGCATCATAGATTTC<br>CATACTTGGATCAGGTTATCATGGAAGCCATGAGATTCTATACTGTGTCTCCATTG<br>ATTGCCAGAGTTACCTCTAGAAGAACTGAATTAGGTGGTCACGAATTGCCAAAAG<br>GTACTTGGTTGTGGATGGCACCTGGTGTTTTATCTAGAGATGCTGCTTCATTTTTT<br>CCAGATCCTGGTGCTTTTAGACCAGAAAGATTTGATCCAGCTTCCGAAGAACAAC<br>GTGGTAGACATCCATGTGCTCATATTCCTTTTGGTATTGGTCCTAGAGCTTGTGTT<br>GGTCAAAGATTTGCCTTGCAAGAATTGAAGTTGTCCATGGTCCACTTGTACCAGA<br>GATTCTTGTTTAGAAGATCCCCACAAATGGAAAGTCCACCAGAATTACAATTCGGT<br>ATCGTCCTGAATTTTAAGAACGGTGTTAAGTTGGTTGCTGTTGAAAGATGTGCTGC<br>TATGTCTTGA |
| *SbMAX1b* | ATGGAAATGGGTACTGTTTTGGGTGCTATGGAAGAGTACACTTTTACTTTTTTGGC<br>TATGGCCGTTGGTTTCTTGGTTTTGGTTTACTTGTATGAGCCATACTGGAAGGTTA<br>GACATGTTCCAGGTCCAGTTCCATTGCCATTGATTGGTCACTTGCATTTGTTGGCT<br>AAACATGGTCCAGATGTTTTTCCAGTTTTGGCCAAGAAACACGGTCCAATTTTTAG<br>ATTCCATGTCGGTAGACAACCATTGATTATAGTTGCTGATGCCGAATTGTGCAAAG<br>AAGTCGGTATTAAGAAATTCAAGTCCATGCCAAACAGGTCTTTGCCATCTCCAATT<br>GCTAATTCCCCAATTCATAGAAAGGGTTTGTTCGCTACTAGAGACTCTAGATGGTC<br>TGCTATGAGAAACGTTATTGTCTCTATCTACCAACCATCTCATTTGGCTGGTTTGA<br>TGCCAACTATGGAATCTTGTATTGAAAGAGCTGCTACCACCAACTTAGGTGATGG<br>TGAAGAAGTTGTTTTCTCCAAGTTGGCTTTGTCTTTGGCCACTGATATTATTGGTC<br>AAGCTGCTTTTGGTACTGACTTTGGTTTGTCTGGTAAACCAGTTGTTCCAGATGAT<br>GATATGAAGGGTGTTGATGTTGTTGTTGGTGATGCTGCTAAAGCTAAAGCTTCTTC<br>TTCCGAATTCATCAACATGCATATCCATTCCACCACCTCATTGAAGATGGATTTGT<br>CAGGTTCTTTGTCTACTATCGTTGGTGCTTTGGTTCCATTCTTGCAAAATCCATTG<br>AGACAGGTTTTGTTGAGAGTTCCAGGTTCTGCTGATAGAGAAATCAATAGAGTTAA<br>CGGTGAGTTGAGAAGAATGGTTGATGGTATCGTTGCTGCTAGAGCTGCAGAAAGA<br>GAAAGAGCACCAGCTGCTACTGCTGCTCAACAACATAAGGATTTTTTGTCCGTTG<br>TTTTGGCTGCCAGAGAATCTGATGCTTCTACAAGAGAATTACTGTCCCCAGATTAT<br>TTGTCTGCTTTGACCTACGAACATTTGATTGCTGGTCCAGCTACTGCAGCTTTTAC<br>ATTGTCATCTGTTGTTTACTTGGTTGCTAAGCACCCAGAAGTTGAAGAAAAGTTGT<br>TAAGAGAAATGGATGCCTTTGGTCCAAGAGGTTCTGTTCCAACTGCTGATGACTT<br>GCAAACTAAGTTTCCTTACTTGGATCAGGTCGTCAAAGAATCTATGAGGTTGTTTA |

| | |
|---|---|
| | TGGTTTCCCCATTGGTTGCAAGAGAAACTTCTGAAAGAGTTGAAATTGGCGGTTA CGTTTTGCCAAAAGGTGCTTGGGTTTGGATGGCTCCAGGTGTTTTAGCAAAAGAT GCTCATAATTTTCCCGATCCAGAGTTGTTTAGACCAGAAAGATTTGATCCAGCTGG TGACGAACAAAAGAAAAGACATCCATACGCTTTCATCCCATTTGGTATTGGTCCTA GAGTATGCATTGGTCAAAAGTTCGCTATCCAAGAAATCAAGTTGGCCATTATCCAC TTGTACCAACACTACGTTTTTAGGCATTCTCCCTCAATGGAATCACCATTGGAATT TCAATTCGGTATCGTCGTTAATTTCAAGCACGGTGTTAAGTTGCACGTTATCAAAA GACACGTTGAGAACAACTAA |
| *SbMAX1c* | ATGGAAATTGCCTTGACTGTTTCCGCTGTTTCTCATCAATCTGTTCCAGTTTTGGT CCTGATCTCTTTCTTGTCTTTGTTCTCTGCTTTCCTGATCTACTTCTATGCTCCATT GTGGTCTGTTAGAAGAGTTCCAGGTCCACCAACTAGATTTCCAATTGGTCACTTG CATTTGTTGGCTAAGAATGGTCCAGATGTTTTCAGAGCTATTGCCAAAGAATACGG TCCAATCTTCAGATTCCATATGGGTAGACAACCATTGGTTATCGTTGCTAATGCTG AATTGTGCAAAGAAGTCGGTATCAAGAAGTTCAAGGACATCAGAAATAGATCTACT CCACCACCATCTATCGGTTCATTGCATCAAGATGCTTTGTTCTTGACTAGAGATTC TACTTGGTCTGCTATGAGATCTACCGTTGTTCCATTATATCAACCAGCTAGATTGG CTGGTTTGATCCCAGTTATGCAATCCTACGTTGATATTTTGGTTGCTAACATTGCT GGTTGGACTGATCAAGATTGCATTCCATTTTGCCAGTTGTCTTTGAGAATGGCCAT CGATATTATTGGTAAGACCGCTTTCGGTATCGAATTCGGTTTGTCTAAAAATGCTG CCGGTGGTGGTGGCGAAACTGAAGGTGGTGAAGGTGATGATAATGTCAGGGAAT TCTTGAAAGAGTACAAGAGGTCTATGGAATTCGTCAAGATGGACTTGTCATCTTCC TTGTCTACTATCTTGGGTTTGTTTTTGCCATGTGTTCAAACTCCATGTAAGAGGTT GTTGAGAAGAGTACCTGGTACTGCTGATTACAAGATGAACGAAAATGAGAGAAGA TTGTGCTCCAGAATCGATGCTATTATTGCTGGTAGAAGAAGAGATAGAGCTACTA GACGTAGAGGTGGTGATGGCGTTTCTGAAGATGATGCTGCTCCATTAGATTTCAT TGCTGCTTTGTTGGATGCTATGGAAAATGGTGGCGGTGCTAAAGATTTTGCTTTG GCTGATAGACATGTTAGAGCTTTGGCTTACGAACATTTGATTGCAGGTACAAAGA CTACCGCTTTCACTTTGTCATCTGTCGTTTACTTGGTTTCTTGCCATCCAAGAGTT GAAGAAAAGTTGTTGAGGGAAGTTGATGGTTTTGCTCCAAGACATGGTAGGGCTC CAGATGCTGATGAATTACAATCAAGATTCCCATACTTGGACCAGGTTATCAAAGAA GCTATGAGGTTCCATTTGGTGTCTCCATTGATTGCTAGACAGACTTCTGAAAGGG TTGAAATTGGTGGTTACGTTTTGCCAAAAGGTGCTTATGTTTGGTTGGCTCCAGGT GTTTTGGCTAGAGATGCTGCACAATTTCCAGATCCAGAAGAATTCAGACCAGAAA GATTTGCTCCTGAAGCTGAAGAAGAAAGAACTAGACATCCATACGCTCATATTCCT TTTGGTGTTGGTCCAAGAGCTTGTATTGGTCATAAGTTCGCTTTACAACAAGTTAA GTTGGCCGTTGTTGAGTTGTACAGAAGATACACTTTTAGACATTCCCCAGCTATG GAATCCCCATTGCAATTTGATTTCGATTTGGTTTTGGCCTTCAGACACGGTGTTAA GTTGAGAGCTATTAGAAGGTCTTAA |
| *SbMAX1d* | ATGGGTTGGGGTGAAATTATCTCTTCCCAGTTGTTGATCGAGTCCTCTTCATCTTC TTTGCCAGCTGTTTTGTTTACTGCTGCTGCTTTGGCTGCTGGTGCATTTGCTGTTT ATTTCTATATTCCATCTTGGAGAGTCAGAAGAGTTCCAGGTCCAGTTGCTTTGCCA TTGGTTGGTCATTTGCCATTATTTGCTAAACATGGTCCAGGTTTGTTCAGGATGTT GGCTAAAGAATATGGTCCAATCTACAGATTCCACATGGGTAGACAACCATTGGTT ATGGTTGCTGATGCTGAATTGTGTAAAGAAGTCGGCATTAAGAAGTTCAAGTCCAT TCCAAACAGATCTATCCCAACTCCAATTAGAGGTTCCCCAATTCATAACAAGGGTT TGTTCTTCACCAGAGACTCTAGATGGCAATCTATGAGAAACGTTATCTTGACCATC TACCAACCATCTCATGTTGCTTCATTGATTCCAGCTATTCAACCATACGTTGAAAG AGCCGGTAGATTATTGCATCCAGGTGAAGAAATTACCTTCTCCGATTTGTCTCTGA AGTTGTTCAATGATACCATTGGTCAAGTTGCCTTCGGTGTTGATTTTGGTTTGACT AAGGATGATACAACTGCTGCTACTTCTCCAGCTGCTCAACAACAACCAGCTCATG GTGGTGCTAATGCTAATCAATCTGTTGATGATCCAGCCACCGATTTCATTAGAAAA CATTTTAGAGCTACCACCAGCCTGAAGATGGATTTGTCTGGTCCATTGTCTATAGT CTTGGGTCAATTTGTTCCATTCCTGCAAGAACCAGTTAGACAGTTGATGTTGAGAG TTCCTGGTTCTGCTGATAGAAGATTGGAAGAGGCTAATTCTGATATGTCTGGTTTG TTGGACGAAATCGTTGCTGAAAGAGCTGCACAAGCTGATAGAGGTCAACAAAAGA ATTTCCTGTCCGTTTTGTTGAACGCTAGAGAATCTACTGAAGCCATGAAGAAGTTG |

| | |
|---|---|
| | TTGACTCCAGATTATGTTTCCGCTTTGACCTACGAACATTTGTTGGCTGGTTCTGT<br>TACTATGTCTTTCACCTTGTCATCCTTGGTTTACTTGGTTGCTATGCATCCAGAAG<br>TCGAAGAAAAGCTGTTGAGAGAAATTGATGCTTTCGGTCCAAAAGATGTTGTTCCA<br>TCTTCTGATGACTTGGAGACTAAGTTCCCATATGTTGAACAAGTCGTCAAAGAAAC<br>CATGAGATTCTATACTGCTTCACCTTTGGTTGCAAGACAAGCTTCTGAAGATGTTG<br>AAGTTGGTGGTTACTTGTTGCCAAAAGGTACTTGGGTTTGGTTGGCTCCAGGTGT<br>TTTGGCAAAAGATCCTAAAGATTTTCCAGATCCAGACGTGTTTAGACCAGAAAGAT<br>TTGATCCTGAATCCGAAGAATGTAAGAGAAGGCATCCATACGCTTTTATTCCATTT<br>GGTATTGGTCCAAGAGCCTGTATTGGTCAAAAATTCGCTATGCAGCAACTGAAATT<br>GGTTGTCATCCACTTGTACCGTAACTACATTTTCAGACATTCCCCAAGAATGGAAT<br>TCCCATTGCAATTCCAATACTCGATCTTGGTCAACTTTAAGTACGGTGTTAAGGTG<br>CAAGTCATCGAGAGAAAGAACTGA |
| *LGS1* | ATGAACGTCCAAGAAAGGCGTAAAGAATTGGAAGAAAGATCTTCTACTACCTTGG<br>GTCACTTGCATACCATTAGAAATACTCCAGCTGGTTCTTCTATGTCTACTACTACTT<br>GTTATTCTGCTCCAGCTGCTGTTGTTCCAGGTGCTGGTGGTGAAGTTGCAGTTGT<br>TACTGCTGTTGCTTCTGAAGCTGGTGCTGCTGCTGCACATGATCAATCTAGAAAA<br>AAGAAGAACAACCACAGGTCCTTGTACGCTAATTTGCCAGCTGCAGAAATTATCG<br>ATTCTTTGCCATTGGAAACTAGGTTCCCAGTTCCACATAGATTATATGGTGGTTTT<br>TGGAAGGCCGAGTTCTTGTTGAAAGGTATGGCTGCAGCTGCTGCTAGAACTACTT<br>CTTGTTTTGAATTCGAGCCAAATCCTTCCGATATTTTCTTGGCTTCATTGCCAAAAT<br>CTGGTACTACTTGGTTGAAGGCTTTGGCTTTTGCTACTTTGAACAGAAGAACTCAT<br>CCACCATCTAATGCTGATGGTCAACATCCATTTTCTCATAGAAACCCACATGACTG<br>CGTTTCGTTCTTGGAATTGATGATGATTCAAGGTGTTGATGCTGCTGCCGCCGAT<br>GACGATGCAGGAGCTCCAAGATTAATCGCGACTCATTTGCCTTGGTCCTGGTTGC<br>CTCCAGCCATAACGGCTGGTGAAGGTCAAGGCGGGGGTTCTTCATCTAGAGGTA<br>GAGGTTGTAGAATCGTTTATGTCTGTAGAGAACCTAAGGACGTCTTGGTTTCTTAC<br>TGGACTTTTTCTGTTAAGGCTGCTGCAAAATTTGCTGCCGCTGCAGCCGCTGGTG<br>GCGACGATGACGGTGGTGGTGGTAGAGAATCTGCTGCAGCTTCTTTGACTACATC<br>TTTTGAAGAAGCTTTCGAGTTGTTCTGCGAAGGTAGATTTCCAGGTGGTCCACATT<br>GGTTGCATGCTTTGGAATTTTGGAGAGAATCTCAAAGACGTCCAGATGAAGTTTT<br>GTTCTTGAGGTACGAAGATATGTTGAGAGATCCAGTTGGTAACTTGAGAAAGTTG<br>GCTGCTTTTATGGGTTGTCCATTCTCTGCTGAAGAAGAAACTGCTGGCGGAGGTG<br>GTGGCGTTGTTGATCAAATAGTTGAATTGTGCTCCTTGGAGAACTTGAAGTCTATG<br>GATGTTAACAAGAACGGTACTACCACTGTTTTGGGTGTTACTAATGATGCCTTTTT<br>CAGAAAAGGTAAGGTTGGTGACTGGAAGAATTACATGACTCCAGATATGGCTGCT<br>AGATTGGATAAGGTTGTCGAAGAAGCTACTAGAGGTTCTGGTTTGACTTTCGCTG<br>ATTCCATTTCTGTCTGA |
| *LGS1-2* | ATGAACGTCCAAGAAAGGCGTAAAGAATTGGAAGAAAGATCTTCTACTACCTTGG<br>GTCACTTGCATACCATTAGAAATACTCCAGCTGGTTCTTCTATGTCTACTACTACTT<br>GTTATTCTGCTCCAGCTGCTGTTGTTCCAGGTGCTGGTGGTGAAGTTGCAGTTGT<br>TACTGCTGTTGCTTCTGAAGCTGGTGCTGCTGCTGCACATGATCAATCTAGAAAA<br>AAGAAGAACAACCACAGGTCCTTGTACGCTAATTTGCCAGCTGCAGAAATTATCG<br>ATTCTTTGCCATTGGAAACTAGGTTCCCAGTTCCACATAGATTATATGGTGGTTTT<br>TGGAAGGCCGAGTTCTTGTTGAAAGGTATGGCTGCAGCTGCTGCTAGAACTACTT<br>CTTGTTTTGAATTCGAGCCAAATCCTTCCGATATTTTCTTGGCTTCATTGCCAAAAT<br>CTGGTACTACTTGGTTGAAGGCTTTGGCTTTTGCTACTTTGAACAGAAGAACTCAT<br>CCACCATCTAATGCTGATGGTCAACATCCATTTTCTCATAGAAACCCACATGACTG<br>CGTTTCGTTCTTGGAATTGATGATGATTCAAGGTGTTGATGCTGCTGCCGCCGAT<br>GACGATGATGCAGATGATGCAGGAGCTCCAAGATTAATCGCGACTCATTTGCCTT<br>GGTCCTGGTTGCCTCCAGCCATAACGGCTGGTGAAGGTCAAGGCGGGGGTTCTT<br>CATCTAGAGGTAGAGGTTGTAGAATCGTTTATGTCTGTAGAGAACCTAAGGACGT<br>CTTGGTTTCTTACTGGACTTTTTCTGTTAAGGCTGCTGCAAAATTTGCTGCCGCTG<br>CAGCCGCTGGTGGCGACGATGACGGTGGTGGTGGTAGAGAATCTGCTGCAGCTT<br>CTTTGACTACATCTTTTGAAGAAGCTTTCGAGTTGTTCTGCGAAGGTAGATTTCCA<br>GGTGGTCCACATTGGTTGCATGCTTTGGAATTTTGGAGAGAATCTCAAAGACGTC<br>CAGATGAAGTTTTGTTCTTGAGGTACGAAGATATGTTGAGAGATCCAGTTGGTAAC |

| | |
|---|---|
| | TTGAGAAAGTTGGCTGCTTTTATGGGTTGTCCATTCTCTGCTGAAGAAGAAACTGC TGGCGGAGGTGGTGGCGTTGTTGATCAAATAGTTGAATTGTGCTCCTTGGAGAAC TTGAAGTCTATGGATGTTAACAAGAACGGTACTACCACTGTTTTGGGTGTTACTAA TGATGCCTTTTTCAGAAAAGGTAAGGTTGGTGACTGGAAGAATTACATGACTCCA GATATGGCTGCTAGATTGGATAAGGTTGTCGAAGAAGCTACTAGAGGTTCTGGTT TGACTTTCGCTGATTCCATTTCTGTCTGA |
| *TaSOT* | ATGAACGCTGCTTTGACTGTTGCTGTTGCTGGTGAAGAAATTGATGAAGCAGTAG CAGGCGAAGAAATAGACGAAGCTGCTTCTAGAGCACAAGCTGATATGTCTGAAAT CATGTCATCTTTGCCAAGATGCCCAGTTTACTTGACTCATCATTATAGAGGTTTCT GGATCAGGGAATTCGTCTTGAAAGGTATGGCTGCTGCTCAAGCTTCTTTTGAACC TAGACCAACTGATGTTTTCTTGGCTTCTTGTCCAAAATCTGGTACTACTTGGTTGA AGGCTTTGGCTTTTGCTACTTTGAATAGAGCTACTCACTTGCCATCTGATTCTAAC CATCCATTGTGTCATAGAAACCCACATGATTGTGTTGCTTTCTTGGAAACTAGACC AGTTCCAGAAACTATGGCTTTGCCATCTCCAAGATTATTGGCTACTCATATCCCAT GTTCTTTGTTGCCATCTAGAATTACCGAATGCGGTAGAGTTGTTTATGTTTGTCCA GAACCTAAGGATGCCTTGGTTTCTTTTTGGATCTACAACAACAAGATCGCCCCAAT GTTGAGAAGAAAGTTTGGTTTGGAATCTCCATCACCAACTTTCGAAGAAGCTTTTG AGTTGTTCTGTGAGGGTCAATCTTCATTTGGTCCACCTTGGAGACATGCTTTGGAA TATTGGGAAGAATCTAGAAGAAGGCCAGGTAAGGTTTTGTTCTTGAGATACGAAG ATATGTTGCAAGATCCAACTGGTAACACTAAGAATTTGGCTGCTTTTATGGGTTGT CCATTCTCTTGTGCTGAAGAAGAAGCTGGCGTTGTTCAAGAAATAGTTCAATTGTG CTCCTTCGAGAAGTTGAAGTCATCTGAAGTTAACAAGAACGGTTCCTCTGCTATGA TGGGTGTTAACAATGATGTCTACTTTAGAAAGGGTGCTGTTGGTGATTGGAAGAA TTATATGACTCCAGAAATGGCTGCCAGGTTGGATAAGATAGTTGAAGAAGCCTTA CAAGGTTCTGGTTTGACTTTCGGTATTTCCATGTGA |
| *ZmSOT* | ATGTACCATCAAACTACCTCCAGACCACAACAAAGACCATCTACTAGACAACCACA TCAAATTCATGCTGTTTTGTTGCCAGATTTGCCAGTTTCTCCAATTGCTTCTACAAG ACGTAGAGCTAGAGGTATTCATTTGTTGCACTTGCAATTGTGCTACTGTAGATCTT GTTGTAACAGAGGTGAAAAGAGAGATCAATTCGCTGGTACTAATACCGCTTGCTG TCATATTTCTGGTATTTTCTTTAGACAAGGCTGGGGTTTGAAAACCGTTGCTGTTT CTTGGGGTTGGGGTTTGTTTCAAGCTGCTGCTTCTAATTGGACTATGATGGCTTCT AGACAGTCTGAAAACAACGCTCAAGAAGAAACTTCTCCATTGACTACTCCAAACG CTAACATTGCCAGAATTATTCCATCTTTGCCATTGGAAACTAGGTGGCCTCCATTT CCATTGAGAAGATATGCTAATTTCTGGTTGCCAGAGGTCACTTTGAAAGAAGGTG TTCCAGGCGTTCATTCTTGTTTTGAACCTAGACCAACTGATGTTTTCTTGGCTTCTT TTCCCAAATCTGGTACTACTTGGTTGAAGGCTTTGGCTTTTGCTACTTTGAAGAGA TCTACTCATCCACCATTCGATGATGATCACCCATTGAGACATTGCAATCCACATGA TTGTGTCAGGTTTTTGGAGTTGGGTTTCAATCAACAAAGGACGAATTGGAAGCTT TGCCATCTCCAAGAGTTTTGGCTACTCATTTGCCATACTCATTATTGCCAGGTTCT ATTACTGGTGATGGTGAACATTCTGGTTGTAGAATCGTTTATGTCTGCAGAGAACC TAAGGATACCTTGGTTTCTTACTGGTTGTTTACTAGAAAAGCTGCTCCAGCTTGTG GTGTTGATGCTAGATCTTTTACTATTCAAGAAGCCTTGGAGTTGTTCTGTGATGGT AGATGTCCAGGTGGTCCACAATGGAATCATGTTTTACAATACTGGAAAGAGTCCG TTAGAAGGCCAGATAGAGTTTTGTTTTTGAGGTACGAAGAAGTCTTGATCGAACCT GAAGCTCATGTTAGAAAGTTGGCTAATTTCATGGGTTGTGGTTTCTCTGAAGAGG AAGAAGAAAGAGGCGTTGTTTCTACTATCGTTGAATTGTGCTCTTTGGGCAAGTTG AGAGATATGGAAGTTAACAGAAACGGTTCTACCAGATTGGGTACTAAGAACGAAT CATTCTTCAGAAAAGGTGTTGCTGGTGATTGGTCTAATCATATGACTCCAGAAATG GCTCACTCCTTGGATAAGGTTGTTGAAGATGCTTTACAAGAGACTGGTTTCACTTT CTCTTCTACCACTTGA |
| *MlSOT* | ATGAGTACAACCACCTGTTATGATTCCGCGACAGCGGTACCTGCAGGAGGAGAG GTGGTTACCGCGGTACCAAGTGAGGAGGCGGCGGCTGCTGCTGTACACCAGAG TAGGAAAAACTTATCCTTGTACGCCAATCTGCCGGCTGCGGAGATCATCGACTCA TTACCGTTGGAAACGCGTTTCCCTGTACCTCACAGGCAATATGGAGGGTTCTGGA AGGCCGAGTTTCTGTTGAAGGGGATGGCGGCTGCAGCAACCAGGTCCACTTGTT |

| | |
|---|---|
| | TCGAACCAAACCCCTCTGACATATTCTTGAGCAGCCTTCCGAAGTCCGGCACGAC<br>CTGGCTTAAGGCCTTGGCATTCGCTACACTGAATCGTGGCACGCATCCACCCTCT<br>AATGCTGACGGGCAACATCCCTTGAGCCATAGAAACCCGCACGACTGCGTTTCAT<br>TCCTGGAGTTAATGATGATTCAGGGCGTTGATGCGGCCGCGGCTGCAGGAGCGT<br>CTGGAGAAGAGCGTGGTTCCCCCCCACCGAGACTAATTGCGACGCACTTACCCT<br>GTTCTTGGCTTCCCCCGGCCATAGTGACGGGCTCAGGATGCAGGATCGTATACG<br>TGTGTCGTGAGCCGAAGGACGTCCTAGTTTCTTATTGGACGTTTTCCGTTAAAGC<br>GGTCGCGAAGTTTGCAGCAGCAGCGGCCGCAGGAGGTGATGGTGACGGGGATG<br>GGGGAGGTGGACGTGAAGCTGCCGCCGCAAGCGCGGGGGGGCTTACTTCATTT<br>GAAGAAGCTTTCGAACTATTCTGTGAGGGAAGATTCCCTGGTGGGCCACACTGGT<br>TGCATGCCCTTGAATATTGGCGTGAGTCTCAGAGGAGGCCGGACGAGGTGCTAT<br>TTTTGAGATACGAGGACATGCTGAGGGACCCAGTAGGGAACTTGAAAAAACTGGC<br>CGCATTCATGGGGTGTCCGTTCTCTGCAGAGGAAGAGAAAGCAGGCGGGGTAGT<br>AGATCAAATCGTAGAGTTGTGTAGCCTGGATAATTTAAGGAGTATGGAAGTCAATA<br>AGAATGGCAGCACAACCGTATTAGGTGTGACGAATGATGCCTTCTTCCGTAAGGG<br>GCAGGTCGGCGACTGGCGTAATTACATGACTCCCGACATGGCTGCTAGATTAGA<br>CAAAGCGGTGGAGGAAGCGACAAGAGGGAGCGGCCTGACGTTCGCAGACAGTA<br>TTGAAGTATAG |
| *ZmMAX1a* | ATGGAAATGGCTGGTGCTGCTGGTACTGAAGCTTGGTTGCCATATGTTACTACTG<br>TTGCTTCTTGTGCTGTTGGCGTTTTTTTCTTGTTGTACTTTTATGCCCCACATTGGA<br>GAGTTAGAGATGTTCCAGGTCCACCAGCTTTGCCAGTTGTTGGTCATTTGCCATT<br>ATTGGCTAGACATGGTCCAGATGTTTTTGGTTTGTTGGCTAAAAAGTACGGTCCAA<br>TCTTCAGATTCCATTTGGGTAGACAACCATTGGTTATAGTTGCTGATCCAGAATTG<br>TGTAGAGAAGTTGGTGTTAGACAGTTCAAGTTGATCCCAAATAGATCTTTGCCAGC<br>TCCAATTGCTGGTTCTCCATTGCATCAAAAGGGTTTGTTTTTCACCAGAGATGAGA<br>GATGGTCTGCTATGAGAAACACCATCATTAGCTTGTACCAACCATCTCATTTGGCT<br>GGTTTGGTTCCAACTATGCAACATTGCATTGAAAGAGCTGCTGACGCTATTCCAG<br>CTATGGTTGTTCAAGAAAATGGTCAGGTTGATTTCTCCGACTTGTCTTTGAAATTG<br>GCCACCGATATTATTGGTCAAGCTGCTTTTGGTGTTGATTTCGGTTTGACTGCTTC<br>TGGTCCAGGTTGTGAAGCTGCTGAATTCATTAGAGAACACGTTCATTCTACCACCT<br>CATTGAAGATGGATTTGTCTGCTCCATTGTCCGTTGTTTTGGGTTTAGTTGCTCCA<br>GCTTTACAAGGTCCAGTTAGACATTTGTTGTCTAGAGTTCCAGGTACTGCTGATTG<br>GAGGGTTGCTAGAACTAATGCTAGATTGAGAGCTAGAGTTGACGAAATCGTTGTT<br>TCTAGAGCAAGAGGTAGAGGTCAACATGGTGAAAGAAGAAAGGATTTCTTGTCTG<br>CTGTTTTGGATGCCAGAGATAGATCTGCTGCTTTGAGAGAATTATTGACCCCAGAT<br>CATGTTTCTGCTTTGACCTACGAACATTTGTTAGCTGGTTCTGCTACTACCGCTTT<br>TACTTTATCTTCAGCCGTTTATTTGGTTGCCGGTCATCCAGAAGTTGAAGCTAAGT<br>TGTTGGCTGAAGTTGATGCATTTGGTCCACATGGTGCTGTTCCAACTGCTGATGA<br>CTTGCAACATAGATTCCCTTATTTGGATCAAGCTTCTGATACCACCATGCAAAGAC<br>ACGTTATCAAAGAAGCTATGAGGTTCTACACTGTGTCTCCATTGATTGCTAGAGTC<br>ACTTCTAGACAAACTGAATTAGGTGGTCATACCTTGCCAAAAGGTACTTGGTTGTG<br>GATGGCTCCAGGTGTTTTATCTAGAGATGCTGCTAATTTTGAAGATCCAGGTGCTT<br>TCAGACCAGAAAGATTTGATCCAGTTTCCGAAGAACAAAGACGTAGACATCCATG<br>TGCTCATATTCCTTTTGGTATTGGTCCAAGAGCTTGTGTTGGTCAAAGATTTGCCT<br>TGCAAGAGGTTAAGTTGAGTATGTTGCACTTGTACAGAAGGTTCTTGTTTAGAAGA<br>TCCCCAAGAATGGAATCACCACCAGAATTACAATTCGGTATCGTCCTGAATTTTAA<br>GAAGGGTGTTAAGTTGGTTGCTGTTGAAAGATGTGCTGCTATGCCATTGTGA |
| *ZmMAX1b* | ATGTTGGCTTCTGCTGTTTTGAGAGCTATGGAAGAATGTACTTTTACCTCTGCTGC<br>TATGGCTGTTGGTTTTTTGTTGGTTGTTTACTTGTACGAGCCATACTGGAAGGTTA<br>GACATGTTCCAGGTCCAGTTCCATTGCCATTTGTTGGTCACTTGCATTTGTTAGCT<br>AGACATGGTCCTGATGTTTTCTTGGTTTTGGCTAAAAAGTACGGTCCAATCTTCAG<br>ATTCCATATGGGTAGACAACCATTGGTTATCGTTGCTAATGCTGAATTGTGCAAAG<br>AAGTCGGCATCAAAAAGTTCAAGTCTATGCCAAATAGGTCCTTGCCATCTGCTATT<br>GCTAATTCCCCAATTCATTTGAAGGGTTTGTTCTCCACTAGAGACTCTAGATGGTC<br>TGCTTTGAGAAACATCATCGTGTCTATCTACCAACCATCTCATTTGGCTGGTTTGA<br>TTCCATCTATGCAATCCCATATTGAAAGAGCTGCTACCAATTTGGATGATGGTGGT |

| | |
|---|---|
| | GAAGCTGAAGTTGCTTTTTCTAAATTGGCTTTGTCTTTGGCCACCGATGTTATTGG<br>TCAAGCTGCTTTTGGTGCTGATTTTGGTTTGACTACAAAACCAGCTGCTCCACCAC<br>CACATCATGGTCCACCAAGACAACATGGTGAAGAGGATGGTGATGGTTCTCATTC<br>TACTAGATCTTCCGAATTCATCAAGATGCATATCCATTCTACCACCTCATTGAAGA<br>TGGATTTGTCTGGTTCTTTGTCTACCATCGTTGGTACTTTGTTGCCAGTTTTACAAT<br>GGCCTTTGAGACAGTTGTTGTTGAGAGTTCCAGGTGCTGCTGATAGAGAAATTCA<br>ACGTGTTAATGGTGCTTTGTGCAGAATGATGGATGGTATTGTCGCAGATAGAGTT<br>GCTGCAAGAGAAAGAGCACCACAAGCTCAAAGACAGAAGGATTTTTTGTCAGTTG<br>TTTTGGCTGCCAGAGATTCTGATGCTGCTGCTAGAAAGTTGTTGACTCCAGATTAT<br>TTGTCCGCTTTGACCTACGAACACTTGTTAGCTGGTTCTGCTACTACTGCTTTTAC<br>TTTGTCATCTGTCTTGTACTTGGTTGCCCAACATCCAAGAGTTGAAGAAAAGTTGT<br>TAAGAGAAGTTGATGCTTTCGGTCCACCTGATAGAGTTCCAACTGCTGAAGATCT<br>ACAATCCAGATTTCCATACACCGACCAAGTCTTGAAAGAATCTATGAGGTTCTTCA<br>TGGTTTCCCCATTGGTTGCTAGAGAAACTTCTGAACAAGTTGATATTGCCGGTTAC<br>GTTTTGCCAAAATCTACTTGGGTTTGGATGGCTCCAGGTGTTTTAGCAAAAGATCC<br>AGTTAATTTTCCAGAGCCAGAGTTGTTTAGACCAGAAAGATTTGATCCAGCTGGTG<br>ATGAACAAAAAAGAAGGCATCCATACGCTTTCATTCCATTTGGTATTGGTCCAAGA<br>ATCTGCATCGGTCAAAGATTCTCTATCCAAGAAATCAAGTTGGCCTTGATCCACTT<br>GTACAGACAATACGTTTTTAGGCACTCTCCCTCTATGGAATCACCATTGGAATTTC<br>AATTCGGTGTCGTCTTGAACTTCAAACACGGTGTTAAGTTGCAGTCCATCAAGAG<br>ACATAAGTGCTGA |
| *ZmMAX1c* | ATGGAAATCACCGCTTCCTGTGATGACGGTGCCGTCACTGCCGGTGCTGTCTCT<br>GGTTTATTGTTGGCTTCTGTCTTGTCTTTGTTCGGTGCTTTCTTGGTTTACTTCTAC<br>GCCCCATTCTGGTCCGTTCGACGTGTTCCAGGTCCTCCTGCCAGATTCCCAATCG<br>GTCATTTGCATTTGCTCGCCAGAAACGGTCCAGATGTCTTCAGAGCTATTGCCAA<br>GGAATACGGTCCAATCTTCAGATTCCACATGGGTAGACAACCATTGGTCATTGTT<br>GCTAATGCTGAATTGTGTAAGGAAGTCGGTATCAAGAAATTCAAGGATATTCCAAA<br>CAGATCAACTCCTCCACCATCTATTGGTTCTTTGCACCAAGATGCTTTATTCTTGA<br>CTAGAGACTCCACCTGGTCTGCTATGAGATCAACTGTCGTTCCATTGTACCAACC<br>AGCTAGATTGGCTGGTTTGATTCCAGTTATGCAATCTTACGTTGACACTCTTGCTG<br>CTAACATTGCTGCTTGTCCAGATCAAGACTGTGTTCCATTCTGCCAATTGTCTTTG<br>AGAATGGCTATTGACATCATCGGTAGAACTGCTTTTGGTATTGAATTTGGTTTATC<br>CAAGAACGCTGCCGGTACTGGTTCCTCTTCTTCTGAATCTCCAGGTGGTGGTGAA<br>GGTGAAGGTGACGTCAGAGAATTCTTAAGAGAGTATAAGAGATCCATGGAATTCG<br>TTAAGATGGATTTGACCTCTTCCTTGTCTACCATCTTGGGTTTGTTTTTGCCATGC<br>GTTCAAACTCCATGTAAGAGACTTTTGAGAAGAGTCCCAGGTACTGCTGACTACA<br>AGATGGACCAAAACGAAAGAAGATTGTGTTCTAGAATTGATGCTATCATTGCTGGT<br>CGCAGACGTGACAGGGCTACCAGAAGAAGATGTGGTCCGGGTGCAGCTCCAGCT<br>CCAGCTCCTTTGGATTTCATCGCTGCTCTTCTGGATGCTATGGAAAGCGGGGGAG<br>GTGGTGGCGGTGGTGCTGGTGCCAACAAGGACTTCGCTCTAGCAGACAGACACG<br>TTAGAGCTTTGGCCTACGAACACTTAATTGCTGGTACCAAGACTACCGCTTTCACC<br>TTGAGTTCTGTTGTGTACTTGGTTTCTTGTCACCCATTGGTAGAAGCTAAGTTGTT<br>GAGGGAATTAGACGGTTTCGCGCCAAGAAGAGGTAGAGGTAGAGCTCCAGATGC<br>TGATGAATTGCAATCCGGTTTTCCATACCTAGACCAAGTTATCAAGGAAGCCATGA<br>GATTCTATGTTGTTTCCCCATTGATCGCTCGTCAAACCTCCGAAAGAGTTGAAATC<br>GGTGGTTACGTTTTGCCAAAGCAAGGTGCTTACGTCTGGTTGGCCCCAGGTGTTT<br>TAGCAAGAGATGCCGCTCAATTCCCAGACCCAGAAGAATTCAGACCAGAAAGATT<br>TGCGCCAGAAGCTGAAGAAGAAAGAGCTCGTCACCCATACGCTCACATCCCATTC<br>GGTGTTGGTCCAAGAGCCTGTATCGGTCACAAGTTCGCCTTGCAACAAGTCAAAT<br>TGGCCGTCGTCGAATTGTACAGAAGATACGTCTTTCGTCATTCTCCATCCATGGA<br>ATCCCCAATTCAATTCGACTTCGACTTAGTCTTAGCTTTCAGACACGGTGTCAAAT<br>TGCGTGCTATCAGAAGAGGTTGA |
| *SbCYP722B* | ATGGATGACATGCACTCTCAATTGCAAGCTGCTGGTGCTGCTTGTCAACAATCTA<br>ATTCTTTGTTGTTGCCACCACCAGCTGCTGATAGACCTTGTTCTTCTTCATCTTCAT<br>CCTCGTTGTCTTTGTTGGGTACAGCTGCTGCTGCATGTTTGTTTTTGTCTGCTGCT<br>ATCTACTGCATCGTCGTTATTATCGTTACTACCTCCTCTAAGCAGAACATCAACAA |

| | |
|---|---|
| | CAGATTGATCAGGCGTTTGTTGAAGTTCAAGGGTAGAAGATCTAAGAACGACAGA AGAAGAGACTACAACAACAATGCTGCTCCACCACCACCACCTCCAGGTAGAGGTT CTTCTTGGTGGTGGTCAGTTGTTGAAACTTTGGCTTTTGTTTCCGCTAACAGATCT GGTAGAGGCTTGTATCATTTCGTTGAAGCTAGACATAGAAGATACGGTCCACCAT GTTTTAGAACTGCTTTGTTAGGTGCTACCCACGTTTTTGTTTCTTCACCAGATGCT GCTAGAAGTTTGTTGGCTGATGCTGGTGGTTTTTCTAAGAGATACGTTAGAACCG TTGCCGAATTATTGGGTGAACATTCTTTATTGTGCGCTTCCCATGATGCTCATAGA GCTTTGAGAAGGGCTGTTGCTCCTTTGTTTAATGCTCAAGCTACTGCTTCTTTGGC TGCTAATTTTGATGCTTTGGCCAGAAGAATTATCACCAGAGATTGGGCTGCTAAAA CTACTGCTGTTGTTGTTTTGGATGCTGCTTTGGATGTTACCTTCGAAGCTATTTGC GATATGTTGATTGGTAGAACTACCACCTTGAAGCGTAGAAGATTACAATCTGATGT TTTGGCTGTTACCAGAGCTATGTTGGCTTTTCCATTGAGATTGCCAGGTACTAGAT TTCATGCTGGTTTGAGAGCCAGAAAAAGAATCATGGATGTCTTGAGACAAGAAAT CGCTTCCAGACAAAGAAACATCATGGATATGGAAGAAATGGAAGAGGATGATTCC AAGCACGACAATGATTTCTTGCAGTCCTTGTTGTTATTGAGGCGTAGAAAGATGAA GTCCTCTCAACAGCAACAATCTCCATCTAACTCTAACGATCATTTGTTCTTGACCG ACGATCAAATCTTGGATAACATCCTGACCTTGATTATTGCTGGTCAAGTTACTACA GCTTCTGCTATTACTTGGATGGTTAAGTACTTGGCCGATAACAAGGATTTCCAAGA AACCTTGAGATCCGTGCAATTGGAAATGGCTTTGAAACACCAACATGGTGATTCT GATGGTCCTTTGACTCTGCAACATTTGAACTCTATGGAATTGGCTTACATGACCGT CAAAGAAAGTTTGAGAATGGCCTCTATCGTTTCCTGGTTTCCAAGAGTTGCTTTGG AAGATTGTCAAGTTGCTGGTTTTCATATCAACAAAGGTTGGATCGTTAACATTGAT GCTAGAGCCTTGCATTATGATGCTACCTTGTATGATAACCCAACCATGTTTGATCC ATCCAGATTCAAAATGGGTGATGGTAGAAGGTGA |
| *SbCYP728B1* | ATGGCTGCTTCTGTTGGTGTTGCTTTGTTGGTTGCTTTTTTGACTCCAGTTGTTGT CTACTTGTTGACCAGACATCCAAACAAAAAACCATTGCCAGGTAATTTGCCACCAG GTTCTTTGGGTTTGCCAATGATTGGTCAATCTTTGGGTCTATTGAGAGCCATGAGA TCTAATACTGGTGAAAGATGGTTGAGAGATAGAGTTGATAGATACGGTCCAGTCT CTAAGTTGTCTTTGTTTGGTGTTCCAACTGTTTTCGTTACTGGTCCAGCTGCTAAC AAATTGGTTTTTGCTTCTGATGCTTTGGCTCCAAAACAACCTAGATGTTTGCCTTT GATTTTGGGCAGAAGAAACATCTTGGAATTGGTTGGTGATGATTACAGACGTGTT AGAGGTGCTATGATGCAATTTTTGAAGCCAGACATGTTGAGAAGATACGTTGGTA CTATTGATGCTGAAGTTGCCAGACACTTGGAAGGTAGATGGGCTGGTAGAAGAAC TGTTGCTGTTTTGCCATTGATGAAGTTGTTGACCTTCGATATTATTGCCACCTTGTT GTTCGGTTTGGAAAGAGGTGCTGTTAGAGAAAGATTGGCTGCTGCTTTTGCTGAT ATGTTAGAAGGTATGTGGTCTGTTCCATTGGATTTGCCATTCACTACTTTCAGAAA GTCCTTGAGAGCTTCTGCTAGAGCTAGAAGAGTTTTGGAAGCTACTTTGGCTGAA AAGAGAGCTAGATTGGAAAGGGGTGAAGCTTCTCCAGCTGATGATTTGGTTTCTT GTTTAGCTTCTTTGAGAGCTGAAGCTGAAGGTGATGGTGGTGAAAGGTTGTTGAC TGATGAAGAAATCGTTGATAACGCCATGGTTGTTTTAGTTGCTGGTCATGATACTT CGTCTGTTTTGATGACCTTCATGATTAGACATTTGGCTGGTGATCCAGCTACATTA GCTGCTATGGTTCAAGAACATGACGAAATTGCTAAGAACAAGGCTGATGGTGAAG CCTTGACTTGGGAAGATCTACATGGTATGAGATTCACTTGGAGAGTTGCTTTGGA AACCTTGAGAATGATTCCACCAATCTTCGGTTCTTTTCGTAGAGCCATGGAAGATA TTGAATTCGATGGTTACTGCATCCCAAAAGGTTGGCAAGTTTTTTGGGCTTCTTCT GTTACTCATATGGACCCATCTATTTTTCCAGATCCAGATAAGTTCCAAGCCTCTAG ATTTGAATCTCAAGCTCCACCATATTCCTTTGTTGCTTTTGGTGCTGGTCAAAGAT TGTGTGCTGGTATTGAATTTGCTAGAGTTGAAACCTTGGTTACCATGCATAGACTA TTGAGAAGGTTTAGATGGCGTTTGTGTTGCGAAGATAAGGATAACACTTTCGTCA GAGATCCAATGCCATCTCCATTGAATGGTTTGCCTATTGAATTGCAGTCTAGAGAT ATGGCTTCTCCAACTCCATCTAAATCTGCTTGTGGTTTGTGA |
| *SbCYP728B35* | *ATGCACATCCCATTGGTTGAAGAATTGAGATTGGCTTCTCCAATGGACTCCTCTTT GATTTTGGCATTGATTTTAGCTGTTGCCTTGGCCTTGTTGTTGCATTTGTTGACAT CTGCTAACAACAAACCTAGAAGGGCTAAACAAGTTCCACCAGGTTCTTTGGGTTT GCCAGTTATTGGTCAATCTTTGTCTTTGTTGAGAGCTATGAGAGCCAATTCTGGTG AAAGATGGATTCAAGATAGAATCCATAGATACGGTCCAGTCTCTAAGTTGTCTTTA* |

| | |
|---|---|
| | *TTTGGTGCTCCAACTGTTTTGTTGGCTGGTCCAGCTGCAAACAAGTTTACATTTTT<br>TTCAAGAGCCTTGGCCATGCAACAGCCAAGATCTGTTCAAAGAATTTTGGGTGAG<br>AAGTCCATCTTGGAATTGGTTGGTGCTGATCATAAGAGAATTAGAGGTGCTTTGG<br>CTGAATTCTTGAAGCCAGATATGTTGAGGTTGTACGTTGGTAAGATTGATGGTGAA<br>GTTAGAAGGCATTTGGACGAAAGATGGGCTGGTAGAACTACTGTTACTGTTATGC<br>CATTGATGAAGAGATTGACCTTCGACATCATCTCGTTGTTGTTGTTCGGTTTACAA<br>AGAGGTGCTCCATTACAAGATGCTTTGGCAGCTGATTTTGCTAGAGTTATGGATG<br>GTATTTGGGCTGTTCCAGTTAATTTGCCATTCACTGCTTTCTCCAGATCTTTGAGA<br>GCTTCTGCTAGAGCTAGAAGATTGATTGCTGGTATTTTGAGAGAAACCAGAGCTA<br>AGTTGGAAACTGGTGAAGCTTCTAGATCCTCTGATTTGATTGCTTGCTTGTTGTCT<br>TTGACCGATCATCATTCTGGTGCACCTTTGTTGTCTGACAAAGAAATCGTTGATAA<br>CTCCGTTGTTGCTTTGGTTGCTGGTCATGATACTTCGTCTATTTTGATGACCTTCA<br>TGGTTAGACAATTGGCCAACGATCCAGATACTTTGGCTGCTATGGTTCAAGAACA<br>TGATGATATCGCTAAGTCCAAAGGTGATGGTGAGGCTTTGGATTGGGAAGATTTG<br>GCTAAAATGAAGTACACTTGGAGAGTCGCTTTGGAAACCTTGAGATTAGTTCCAC<br>CTATGTTTGGTGATTTTAGAAGGGCATTGCAAGACGTTGAATTCGATGGTTACTTG<br>ATTCCAAAAGGTTGGCAAGTTTTTTGGGTTGCTTCTGTTACTCATATGGATCCAGG<br>TATTTTTCCAGAACCAGCTAGATTTGAACCATCCAGATTCGAAAATCAATCCCCAC<br>CATGTTCATTCGTTGCTTTTGGTGGTGGTCCAAGAATTTGTGTTGGTATGGAATTC<br>GCTAGAATCGAAACTTTGGTTACCATGCACTATTTGGTGAGAAGATTCAGATGGAA<br>GTTGTGCTGTAAGAAGGATACTTACGCTAGAGATCCAATGCCATTGCCATTGCAT<br>GGTTTGCCAATTCAATTGGAACATAAGGTTTCTCCATGCGTCATGTGA* |
| *ZmCYP728B3<br>5* | *ATGGACTCCTCTTTGGTTTTGGCTTTGATTGCTGTTGCTTTGCCAGTTTTGTTGCA<br>CTTGTTGAAAAGAGGTAAAACTCCTTGGAGGCCAGCTGCTGCTAAATTGCCACCA<br>GGTTCTTTGGGTTTACCAGTTATTGGTCAATCCATCGGTTTGTTGAGAGCTATGAG<br>AGCAAATACTGCTGAAAGATGGATCTTGGATAGAATCCATAGATACGGTCCAGTC<br>TCTAAGTTGTCTTTGTTTGGTAGACCAACTGTTTTGGTTGCTGGTTCAGCTGCTAA<br>TAGGTTCATTTTTTTCTCATCCGCTTTGGCTATGCAACAGCCAAGATCTGTTCAAA<br>GAATTTTGGGTGACAAGTCCATCTTGGAATTGACTGGTGCTGATCATAAGAGAATT<br>AGAGGTGCTTTGGTCGAATTCTTGAAGCCAGATATGTTGAGGTTGTACGTTGGTA<br>AGATTGATGGTGAAGTTAGAAGGCATTTGGATGAATGTTGGGCTGGTAGATGTAC<br>TGTTACTGTTATGCCACATATGAAGAGATTGACCTTCGACATCATCAGCTTGTTGT<br>TGTTTGGTTTGGAAAGATCCCCATTGCAAGATGCTTTGGCTGGTGATTTTGCTAGA<br>GTTATGGATGGTATTTGGGCTGTTCCAGTTAATTTGCCATTCACTGCTTTCTCCAG<br>ATCTTTGAGAGCTTCTGCTAGAGCTAGAAGATTGATTGCAGGTATTGCTAGAGAAA<br>CCAGAGCTAAATTGGAAAGAGGTGAAGCTTCTAGATCCTCTGATTTGATAGCATG<br>CTTGTTGTCCTTGACTGATCATTCTGGTGCTAGATTGCTATCCGAAGAAGAAATCG<br>TTGATAACTCCATGGTTGCTTTAGTTGCTGGTCATGATACCTCTTCTATTCTGATG<br>ACTTTCATGGTTAGACACTTGGCTAATGATCCAGATACTTTAGCTGCTATGGTTCA<br>AGAACATGACGAAATCGCTAAGAACAAAGGTGATGGTCAAACTTTGGATTGGGAA<br>GATTTGGCTAAGATGAAGTACACTTGGAGAGTTGCTTTGGAAACCTTGAGATTGG<br>TTCCACCAATTTTTGGTAACTTTAGAAGGGCCATGCAAGATATTGAATTCGACGGT<br>TACTTGATCCCAAAAGGTTGGCAAGTTTTTTGGGCTGCTTCTGTTACTCATATGGA<br>TACTGGTATTTTCCATGAACCAGCTAAGTTCGATCCATCCAGATTCGAAAATCAAT<br>CTGCTGCTTCAGCTCCACCATGTTCATTTGTTGCTTTTGGTGGTGGTCCAAGAATT<br>TGTGTTGGTATGGAATTCGCTAGAATCGAAACTTTGGTTACCATGCACTATTTGGT<br>GAGAAGATTCAGATGGAAGTTGTGCTGTAAGAACGATACTTTCGCTAGAGATCCA<br>ATGCCATCTCCATTGCATGGTTTGCCAATTGAATTGGAACAAAAGGCTTCTCCCTG<br>A* |

**Table S7.** Primers used for site-directed mutagenesis of LGS1

| Primer name | Sequence (5'-3') |
|---|---|
| H216A-F | TCGCGACTGCTTTGCCTTGGTCCTGGTTGCC |
| H216A-R | AAGGCAAAGCAGTCGCGATTAATCTTGGAG |
| H317A-F | CATTGGTTGGCTGCTTTGGAATTTTGGAGAG |
| H317A-R | CAAAGCAGCCAACCAATGTGGACCACCTGG |
| Y247F-F | GAATCGTTTTTGTCTGTAGAGAACCTAAGGA |
| Y247F-R | TACAGACAAAAACGATTCTACAACCTCTAC |
| K148A-F | CATTGCCAGCTTCTGGTACTACTTGGTTGAA |
| K148A-R | TACCAGAAGCTGGCAATGAAGCCAAGAAAA |
